# Long-range action of an HDAC inhibitor treats chronic pain in a spared nerve injury rat model

**DOI:** 10.1101/2023.12.13.571583

**Authors:** Maria Virginia Centeno, Md. Suhail Alam, Kasturi Haldar, Apkar Vania Apkarian

## Abstract

Histone deacetylase inhibitors (HDACi) that modulate epigenetic regulation and are approved for treating rare cancers have, in disease models, also been shown to mitigate neurological conditions, including chronic pain. They are of interest as non-opioid treatments, but achieving long-term efficacy with limited dosing has remained elusive. Here we utilize a triple combination formulation (TCF) comprised of a pan-HDACi vorinostat (Vo at its FDA-approved daily dose of 50mg/Kg), the caging agent 2-hydroxypropyl-β-cyclodextrin (HPBCD) and polyethylene glycol (PEG) known to boost plasma and brain exposure and efficacy of Vo in mice and rats, of various ages, spared nerve injury (SNI) model of chronic neuropathic pain. Administration of the TCF (but not HPBCD and PEG) decreased mechanical allodynia for 4 weeks without antagonizing weight, anxiety, or mobility. This was achieved at less than 1% of the total dose of Vo approved for 4 weeks of tumor treatment and associated with decreased levels of major inflammatory markers and microglia in ipsilateral (but not contralateral) spinal cord regions. A single TCF injection was sufficient for 3-4 weeks of efficacy: this was mirrored in repeat injections, specific for the injured paw and not seen on sham treatment. Pharmacodynamics in an SNI mouse model suggested pain relief was sustained for days to weeks after Vo elimination. Doubling Vo in a single TCF injection proved effectiveness was limited to male rats, where the response amplitude tripled and remained effective for > 2 months, an efficacy that outperforms all currently available chronic pain pharmacotherapies. Together, these data suggest that through pharmacological modulation of Vo, the TCF enables single-dose effectiveness with extended action, reduces long-term HDACi dosage, and presents excellent potential to develop as a non-opioid treatment option for chronic pain.

## Introduction

Chronic pain is a leading source of disability worldwide (*1, 2*). Recent estimates suggest that in the US alone, more than 51 million people, greater than 20% of adults, suffer chronic pain(*3*) Its incidence is higher than many other prevalent disorders, and frequently, its persistence is long-term, lasting years to a lifetime (*4*). Yet treatment options remain limited: available medicines result in short-lasting pain relief, with significant adverse effects, and the most effective options, namely opioids, are linked to addiction and related mortality at epidemic rates (*5, 6*). Thus, chronic pain presents a high unmet clinical need where novel, non-opioidergic treatment options are urgently required (*7–9*).

Chronic pain is most often a persistent state arising from a prior acute injury, suggesting involvement of epigenetic mechanisms (*10*). Histone deacetylases (HDACs) are key players in epigenetic regulation (*11*) and their inhibitors (HDACi) have progressed rapidly in the treatment of cancers (*12*). But due to the acute need for treatments, and since epigenetic mechanisms are important for nervous system function at all stages (*13–15*), HDACi are also broadly of interest for their potential to treat a wide range of neurological diseases (*16–20*), including chronic pain (*21*) In tumor killing, HDACi also targets immune cells in the tumor microenvironment (*22–25*). The drugs act by blocking one or more classes of HDAC enzymes to induce acetylation of histones and promote the opening of chromatin (*26*) to remodel innate immune cell macrophage and myeloid cell function, which in turn further modulate adaptive immune responses (*27, 28*). Both innate and adaptive immune cell responses are known to underlie inflammatory pain, and as summarized in recent reviews (*21, 29–31*), multiple studies have reported that HDACi reduce inflammatory responses and modulate chronic pain (as measured by changes in allodynia and hyperalgesia) in a range of animal disease models (*32–34*).

Major challenges in utilizing HDACi to treat chronic pain arise because most HDACi poorly permeate the central nervous system (CNS) and target multiple HDACs. HDACs encode essential functions (*35–37*) and hence, the extent of their blocking that can be tolerated for short-term tumor treatment (over 4 weeks), may not be acceptable for months to years-long treatments expected for chronic pain. Although HDACi designed to have better penetration of the central nervous system and greater selectivity are emerging (*38–41*), the five FDA-approved HDACi remain broad spectrum and largely excluded from the CNS. However, since re-purposed drugs have been shown to have a much shorter time to clinic than new chemical entities (*42*), we investigated pharmacological approaches with the first FDA-approved HDACi, Vo, to improve long-term treatment of a monogenetic neurological disease in mice (*43*). This was accomplished through the development of a triple combination formulation (TCF) comprised of Vo (at its FDA-approved daily dose), 2-hydroxypropyl-β-cyclodextrin (HPBCD), and polyethylene glycol (PEG) that boosted plasma and brain exposure and efficacy of Vo. We showed that the TCF administered weekly protected against neurodegeneration and neuroinflammation at ∼14% of the total weekly Vo dose approved to treat tumors (*43*). Importantly, despite its improved exposure, Vo in TCF was nonetheless eliminated from the brain and body within four hours (compared to Vo alone, which was eliminated in 2h)(*43*). Thus, TCF-treated animals were free of Vo and could resume HDAC for over 97% of the time between weekly injections. Indeed, long-term administration of weekly TCF over 8-10 months in mice failed to impair brain or neuromuscular functions based on quantitative analyses of neurons, neuroinflammation, neurocognitive/muscular disease symptom progression, cerebellar/hippocampal nerve fiber-staining, and *Hdac* gene-expression (*44*).

Here we assessed the efficacy of TCF and its components in a spared nerve injury (SNI) rat model of neuropathic chronic pain. Our results show that less than 1% of the total monthly Vo anti-tumor dose, when delivered via a single injection of the TCF, significantly reduced allodynia over a period of four weeks. Doubling Vo in TCF, tripled the response amplitude and alleviated allodynia over many months. These data strongly support that through pharmacokinetic modulation, the TCF confers effectiveness to a single dose of Vo with long-term action. Comparative analyses in the mouse SNI model, suggests the action is sustained days/weeks after the drug is eliminated from the animals. This substantially reduces total dosing over a significant treatment period: in summary, the TCF presents the best-in-class potential to further explore as a non-opioid treatment option for chronic pain.

## Materials and Methods

### Materials

All fine chemicals were purchased from Sigma (St Louis, MO, USA) unless otherwise indicated. Vorinostat was procured from Selleck Chemicals (Houston, TX, USA).

### Animals

For these experiments, we used rats and mice. Except for the first experiment in which we used Sprague-Dawley aging male rats, the rest of the rats used were Sprague-Dawley weighting between 200 to 250 grams. We also used adult male C57BL/6 mice weighing 25 to 31 grams. The animals were group-housed and had free access to standard food and water. The described experiments were performed at Northwestern University, Chicago, and at Boler-Parseghian Center for Rare and Neglected Diseases, Department of Biological Sciences, University of Notre Dame, Indiana. The animals were kept at 21 ± 2°C temperature and 30% to 60% humidity, under a 12/12-hour light/dark cycle. Handling and testing were performed during the light period. To minimize stress, they were handled regularly before surgery and behavioral testing. All studies were approved by the Animal Care and Use Committee of Northwestern University as well as the University of Notre Dame, followed the ethical guidelines for the investigation of experimental pain in conscious animals.

### Neuropathic pain model: SNI and Sham

The SNI model has been described previously (*45*). Animals were anesthetized with isoflurane 1.5–2% and a mixture of 30% N2O and 70% O2. The sciatic nerve was exposed at the level of trifurcation into the sural, tibial, and common peroneal nerves. The tibial and common peroneal nerves were tightly ligated and severed, leaving the sural nerve intact. Animals in the Sham surgery group had their sciatic nerve exposed as in the SNI procedure but received no further manipulation.

### Drug injections and organ harvest

The triple combination formulation (TCF) was a mixture of vorinostat (50mg/Kg), 2 hydroxypropyl-*β*-cyclodextrin (HPBCD, 2000mg/Kg), polyethylene glycol 400 (PEG, 45%) and DMSO (5%). Vehicle contained HPBCD (2000 mg/Kg), PEG 400 (45%), and DMSO (5%). Vorinostat (50 mg/Kg) was prepared in 5% DMSO and 45% PEG. Detailed methodology for preparing TCF drug solutions has been described earlier (*43, 44*). Injection volume was 5.6 ml/kg in rats and 10 ml/kg in mice. With the exception of the data in Fig. 1 A (LHS panel) all animals received 2% lidocaine, 5 to 10 minutes before drug and or vehicle treatment. TCF or its components were administered weekly through the intraperitoneal route into rats of specified ages and frequency. 2xTCF was prepared using Vo (100 mg/Kg) in 5%DMSO and 45% PEG. Rats were anesthetized using ketamine/xylazine (80 and 10 mg/0.1 kg, respectively) and, perfused transcardially with saline followed by 4% paraformaldehyde. The spinal cord was isolated, flash-frozen in liquid nitrogen, and stored at −80 °C or immersion fixed in 10% neutral buffered formalin.

**Figure 1.**
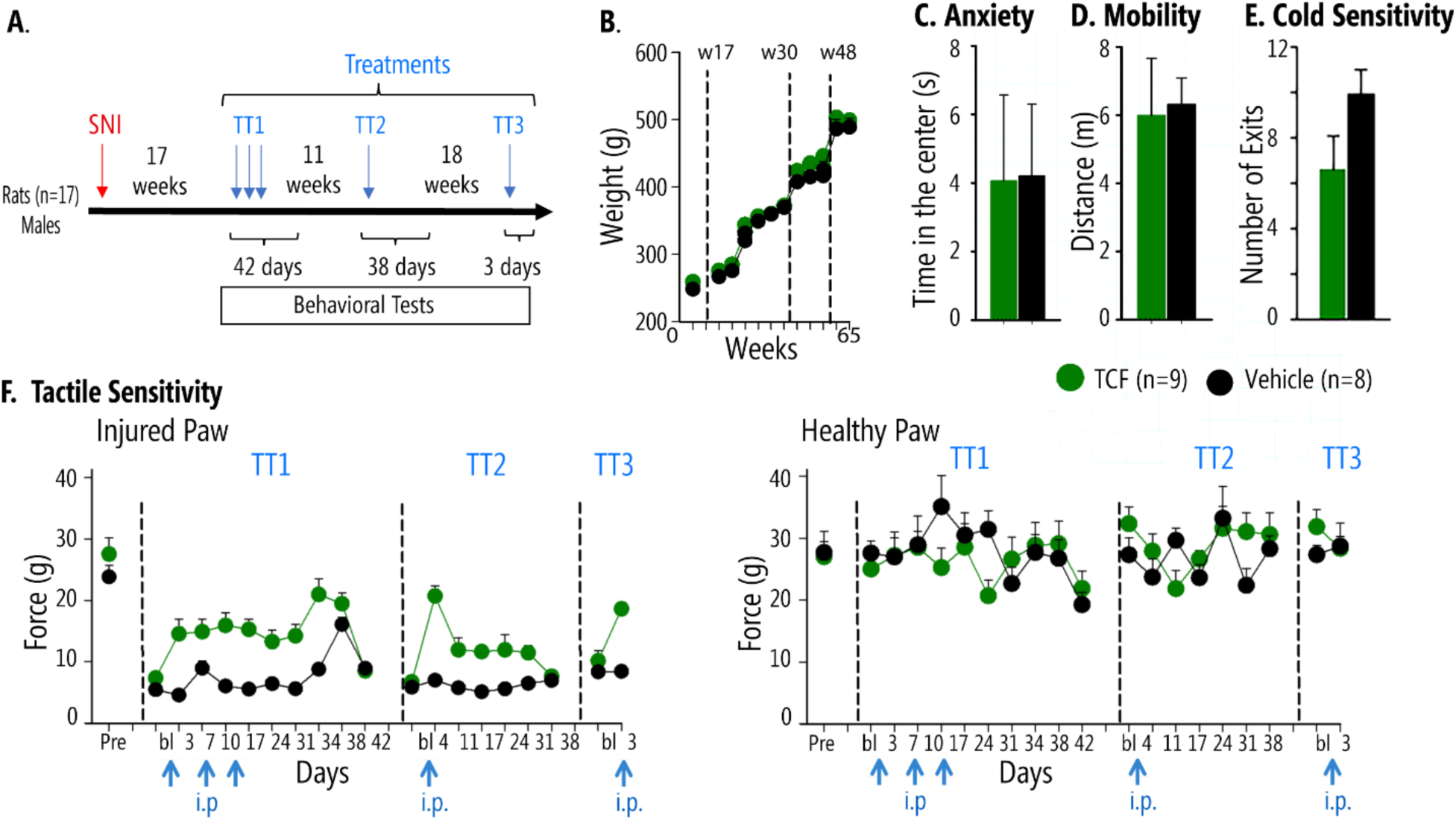
Effects of TCF on behavioral measures in chronic neuropathic (SNI) male rats, 17 weeks after peripheral injury. **A.** Timeline and schedule of TCF administration (blue arrows). Male rats underwent SNI surgery, 17 weeks later behavior was assessed for TCF (n = 9) or vehicle (n = 8) administrations. **B.** Weight increase over 65 weeks (spanning TT1-TT3 treatments) was not different between TCF-treated and vehicle-treated SNI rats. **C&D.** Anxiety-like behavior and mobility tested 3 days after TT3 were not different between TCF-treated and vehicle SNI mice. **E.** There was an improvement in cold sensitivity tested 4 days after TT3 in TCF-treated SNI relative to vehicle, as assessed by cold surface place avoidance (number of exits from cold surface chamber, p=0.028). **F.** Pain-like behavior as assayed by tactile sensitivity (tactile allodynia, smaller values indicate more sensitivity implying more pain). TCF, but not vehicle, diminished pain-like behavior only for the SNI paw, for long periods, reversibly and reproducibly. Left panel, SNI injured paw, pre-injury baseline (Pre) remains decreased 17 weeks after SNI (bl). Left panel, TT1: A single injection of TCF, in contrast to vehicle, statistically significantly relieves tactile allodynia. Weekly application of TCF or vehicle shows the same extent of pain relief, which remained significantly different from vehicle for two-weeks after the last TCF injection (2-way repeated measures ANOVA, F_10,150_=5.1 p<0.001 for treatment*time, See Supp. Table 1 panel A). By day 42, TCF effects dissipated, and tactile allodynia was not different from vehicle. Left panel, TT2: Tactile sensitivity at baseline (bl) was not different between treatments, but a single TCF injection, but not vehicle, (11 weeks after the last TT1 injection) resulted in pain relief for 30 days (2-way-ANOVA F_6,90_=6.878 p<0.001 for treatment*time, Supp. Table 1 panel B). Left panel, TT3: At week 48, 18 weeks after TT2, TCF but not vehicle injection again decreased pain-like behavior (2-way-ANOVA F_1,14_=15.77, p=0.001 for treatment*time, Supp. Table 1 panel C). Right panel, TT1-TT3: no differences were seen in tactile allodynia between TCF and vehicle in the uninjured paw. Numbers are mean±SEMs.

### RNA extraction and qPCR analysis

Total RNA from the frozen ipsilateral side of the spinal cord was extracted using RNeasy Plus Universal kit from Qiagen and quantitative PCR (qPCR) was performed using Power SYBR Green RNA-to-CT 1-step Kit. Sequences of primers were as following. *Gfap, forward 5’-*GGTGGAGAGGGACAATCTCA-3’, *reverse, 5’-*CCAGCTGCTCCTGGAGTTCT-3’ *Cd11b, forward, 5’-* CTGGGAGATGTGAATGGAG-3’, *reverse,* 5’-ACTGATGCTGGCTACTGATG-3’ *Iba1, forward, 5’-* GCCTCATCGTCATCTCCCCA-3’, *reverse,5’-* AGGAAGTGCTTGTTGATCCCA-3’ *and Gapdh, forward, 5’-*ATGACTCTACCCACGGCAAG-3’ *and reverse, 5’-*TACTCAGCACCAGCATCACC-3’ GAPDH was used as endogenous control. The relative quantification of gene expression was done by the *ΔΔ*C_T_ method.

### Von Frey Assay

Paw withdrawal thresholds to von Frey filament stimulation (VF) were used to assess mechanical sensitivity of the hind paws. Animals were placed in a Plexiglas box with a wire grid floor and allowed to habituate for 15 min for rats and at least 50 minutes for mice. At this point, filaments of various forces (Stoelting) were applied to the plantar surface of each hind paw. Filaments were applied in a descending or ascending pattern, determined by the response of the animal. Each filament was applied for a maximum of 2 seconds, and paw withdrawal in response to the filament was considered a positive response. Fifty percent thresholds were calculated following the method described in (*46*). A lower 50% threshold value suggests higher tactile sensitivity, implying more severe pain-like behavior.

### Cold sensitivity (acetone test)

A blunt needle connected to a syringe was used to drop 50 μL of acetone on the lateral hind paw. Mice were observed for 5 minutes, and their withdrawal behavior and the duration of their withdrawal reaction were recorded according to previous reports (*47*). Paw withdrawals due to locomotion or weight shifting were not counted.

### Cold sensitivity (place aversion test)

Animals were placed in an arena (66cm x 12cm x18cm) divided into two equal size hemi-chambers to which they had free access. The floor of one of the hemi-chambers was at neutral room temperature with bright light and the other was at 10C (cooled by TECA Compact Air Cooled Thermoelectric Cold Plate) with dim light. The activity was measured for 15 minutes. Any-maze automated video recording and software system (v4.99, Stoelting Co.) quantified the total time spent in each hemi-chamber and the amount of exits from the cold zone.

### Open field

Animals were placed in a plexiglass box (60cm × 60cm x 50 cm for rats and 50cm x 50cm x 30cm for mice) under dim light. Locomotor activity (distance traveled in the box indicative of general level of activity) and anxiety (time spent in the center of the field and in the periphery of the field) were recorded for 5 minutes and analyzed using the video ANY-maze Visual Tracking System and Software (v4.99, Stoelting Co.).

### Novel Object recognition

This test was originally described by Ennaceur and Delacour (*48*). Behavior is evaluated in a plexiglass box (50×50×30 cm) under dim light. The test is run in two days and consists of three steps: Habituation, Training, and Test. For habituation, the rat is placed in the empty test box and allowed to explore the space for 15 minutes on two consecutive days. The Training session starts 10 minutes after the second habituation session. The rat is allowed to visually explore two identical objects for 10 minutes. One hour later, the rat is placed back in the testing box where one of the previously explored objects is replaced with a novel object, and then, the rat is allowed to explore the new pair of objects for 10 minutes. The objects used were made of plastic, preventing the animals from easily displacing them during exploration. A top camera monitors the time the rat spends with each object. The box and the objects were cleaned with a 70% ethanol solution after each trial. Because rodents have an innate preference for novelty, a rodent that remembers the familiar object will spend more time exploring the novel object.

### Immunohistochemistry, image acquisition and analysis

Tissue embedding and immunohistochemical staining of spinal cord sections were performed by NeuroScience Associates (Knoxville, TN, USA). Thirty-micron thick sections were cut through the entire depth of the ipsilateral side of vehicle (Veh) and TCF-treated and contralateral side of a vehicle and TCF-treated animals, respectively. Free-floating sections after blocking were incubated in rabbit anti IBA-1 antibody (Abcam, ab178846, 1:75000) for overnight at room temperature. Biotinylated secondary antibodies were from Vector Laboratories (BA-1000, 1:1000). Sections were developed following standard methods. To quantify microglial abundance IBA1 stained spinal cord slices from 7 depths at the interval of 180µm were analyzed and bright field images were captured using A10 PL 10x (for counting) objective lens (Nikon) with a numeric capture of 0.25. After applying a constant threshold, numbers were automatically counted using ImageJ. Numbers are mean±SEM. P value as indicated by Student’s t-test. ns, non-significant.

### Statistical Test

Graphs were plotted in MS Excel or GraphPad Prism 8.4.3. For comparing two groups student’s *t-test* or Mann-Whitney U test was performed. For comparing more than two groups, one-way ANOVA with Tukey’s post hoc analysis (parametric) or Kruskal-Wallis test (nonparametric) was performed. All statistical tests were done with GraphPad Prism 8.4.3 and were two-tailed. Statistical tests and P values are indicated in the figure legends. *P<0.05* was considered significant. *\*P<0.05, **P<0.01, ***P<0.001, ****P<0.0001*.

## Results

### Effects of the TCF on chronic pain and spinal cord inflammation in the SNI rat model

Prior studies have suggested the levels of SNI-induced mechanical allodynia are highest in aged rats (*49, 50*) that are often refractory to treatments (*50*). We, therefore, initiated testing the effects of TCF in adult Sprague-Dawley rats 17 weeks (day 119) after SNI (a model for long-term chronic neuropathic pain). After the baseline response was tested, we injected a single dose of TCF in 9 male SNI rats, while a control group of 8 males received vehicle (which comprised of HPBCD, PEG and DMSO but lacked Vo) (Figure 1A shows the timeline and experimental design). As shown in Figure 1 (F, left panel), by day 3, there was a significant reduction in mechanical allodynia in rats treated with TCF-compared to vehicle. Since in our prior studies on protecting neurodegeneration, TCF was administered weekly; we repeated injections 7 and 14 days after the first injection to find that reduction in allodynia continued to be observed well after a month of the initial injection, while repeat TCF injections did not increase pain relief effect size. Tactile sensitivity showed no changes when tested on the uninjured paw (Figure 1F right panel, TT1). These and additional behavioral assays, the cold plate, open field, and acetone tests were continued up to 42 days after the first injection after which the animals were allowed to rest, till allodynia reached baseline. Our data suggested that TCF reduced mechanical allodynia in the SNI rat model. Moreover, the benefits of a TCF injection to mechanical allodynia may last beyond a week.

To test the length-of-time benefit from a single injection, the same rats that returned to baseline allodynia at week 30 (day 210) were given a single injection of the TCF or vehicle. As shown in Figure 1F, left panel, TT2, TCF specifically reduced allodynia (no effect on uninjured paw, Figure 1F, left panel, TT2) for over a month as measured in behavioral assays over 38 days, but without effect on weight, anxiety or mobility (Figure 1B-D) and a decreased sensitivity to cold (Figure 1E, group difference for exits from cold plate p = 0.028). Together these data strongly suggested that TCF may have a long-lasting analgesic effect on mechanical allodynia mediated by large fibers, but a smaller effect on sensitivities caused by small fibers and had no side effects on tactile sensitivity for the uninjured paw, weight, anxiety, and mobility.

Chronic pain is strongly associated with inflammatory changes (*21, 51*). In the SNI model, prominent changes are seen in the spinal cord (*51*). We, therefore, evaluated the effects on inflammation in the spinal cord of SNI rats as a consequence of TCF or vehicle treatment. To do this, we injected animals that had returned to baseline allodynia at 48 weeks (day 336; Figure 1A, TT3). After the reduction of allodynia was observed at day 3 (Figure 1F, left panel, TT3), the animals were sacrificed, spinal cord was harvested and processed for both immunohistochemical and molecular analyses. As shown in Figure 2A, when we probed spinal cord sections for Iba1 protein by immunohistochemistry, we detected its location in microglia in sections from both ipsilateral (Fig. 2A upper panels) and contralateral (Fig 2A, lower panels) spinal cord regions in vehicle and TCF treated animals. Quantitative analyses (Fig. 2B) suggested elevation of microglia on the ipsilateral side seen with vehicle treatment (and a subset of which were larger, suggesting activation; see yellow arrowheads, Fig. 2A, inset, LHS upper panel), that was effectively reduced and neutralized by TCF (Fig. 2B and Fig. 2C). On the contralateral side microglia levels were comparable between TCF and Vehicle and both were reduced (Fig. 2B). In contrast qPCR analyses failed to show a reduction in transcript levels of Iba1 (Fig. 2C). However, TCF did reduce transcript levels of Cd11b, a marker for activated microglia, relative to vehicle (Fig. 2 D). Together, these data suggest there was activated microglial increase associated with sidedness of injury in SNI animals receiving vehicle (which is consistent with prior studies showing that these immune cells are associated with spinal cord inflammation (*52*)): and TCF reduced microglial levels to baseline. TCF also reduced transcript levels of Gfap (Fig. 2E, a marker for astrocytes, which are also a major glial cell type), consistent with the hypothesis that a mechanism of action of the TCF may be through the reduction of inflammation.

**Figure 2.**
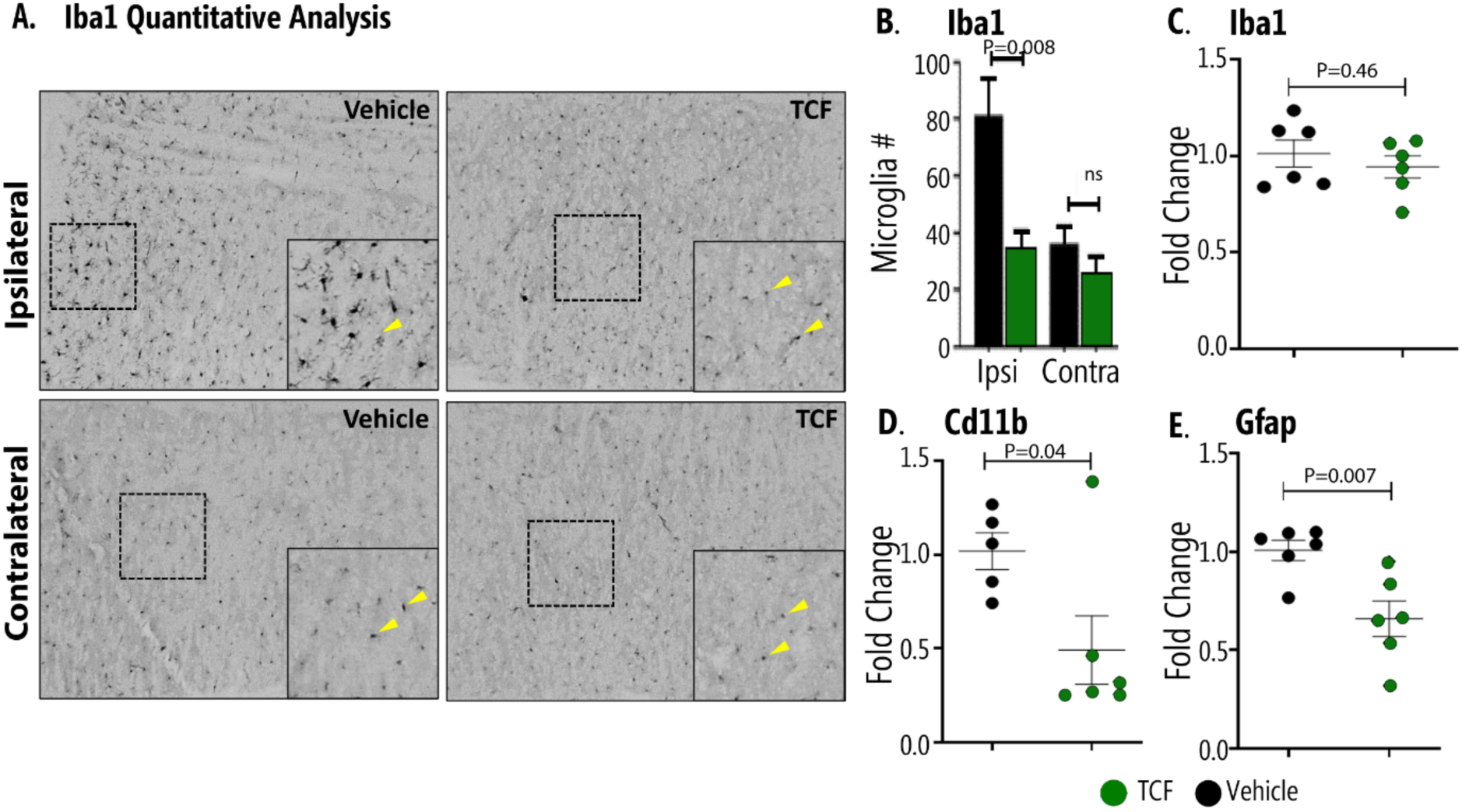
Effects of TCF on spinal cord inflammatory responses of SNI rats. **A.** Immunohistochemical analysis of the distribution of microglia (marked by the inflammatory protein Iba1) of the spinal cord of SNI rats. Representative images show Iba1 positive microglia (yellow arrow), in the ipsilateral and contralateral sides of vehicle and TCF treated animals. Insets are zoomed region shown by dotted box. **B.** Quantitative analysis of the data in A taken from seven different depths**. C-E.** Fold change in transcripts of (**C**) Iba1, (**D**) Cd11b and (**E**) Gfap. Control: GAPDH. N=5-6 per rat group. Mean± SEM are shown. P values as indicated, student’s t test. Numbers are mean±SEMs.

### Action of a single dose of TCF in reducing long-acting allodynia is mirrored in repeat injection

To confirm that a single injection of TCF is necessary and sufficient to reduce allodynia, we compared effects of TCF (n=7), Vo (n=5), Veh (n=6) and a control of no injection (∼TT, n=6) on the injured paw in a total of 24 male SNI rats at 2 months of age, ten days after SNI, that had no prior exposure to drugs (see Figure 3A for experimental design and timeline). As shown in Fig. 3B (left panel, early treatment) after a single injection, the TCF significantly reduced allodynia with a measurable response seen on day 1, that peaked by week 1, was lowered in week 2 but nonetheless sustained to between 3-4 weeks. Vo and Vehicle tactile allodynia responses remained at the baseline of untreated animals throughout this period. Two weeks after the rats reached baseline (or 55 days after SNI), a repeat TCF injection was administered (Fig. 3A, second injection) with the control groups receiving Vehicle, Vo, or no treatment. As shown in Fig. 3B right panel, TCF rapidly reduced allodynia by day 4, with sustained activity up to week 3. Vo in PEG also showed a response but began to approach baseline by week 3, when levels of allodynia reduction sustained by TCF were significantly greater (which was also seen earlier in the treatment period). Vehicle and no treated groups of animals effectively kept the response at baseline. Examination of the healthy paw, shown in Fig. 3C, showed that in both early and later treatment regimens, TCF, Vo, Vehicle, or saline had no effect on allodynia, establishing that drug action is limited to the injured paw.

**Figure 3.**
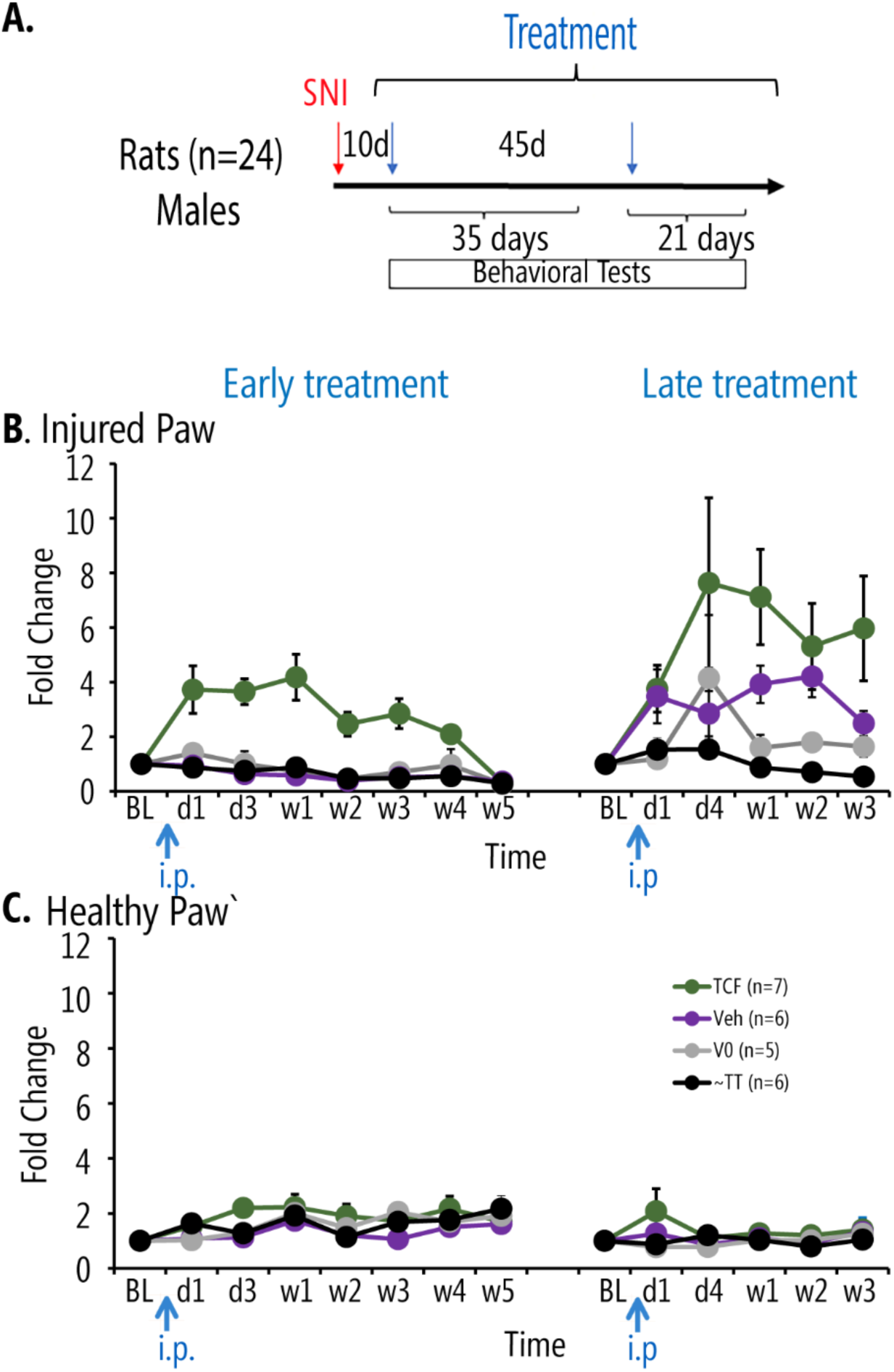
Effects of single TCF injections on tactile allodynia in young SNI male rats. **A.** Timeline and schedule of behavioral assessments (red arrow=SNI injury; blue arrows=TCF or control injections). **B**. Change in tactile allodynia during early (2-way, repeated-measures ANOVA 2-RM-ANOVA F_21,140_=5.563, p<0.0001 for drug* time) and late (2-way, repeated-measures ANOVA 2-RM-ANOVA F_15,100_= 1.673, p=0.068 for drug* time) treatments for the SNI injured paw. **C.** Change in tactile allodynia for the uninjured paw (n.s.). Numbers are mean±SEMs.

In these animals we also tracked cold sensitivity (using acetone test) and mobility (open field test) over time (see Supp. Figure 3). There were small disturbances in these measures only one hour after TCF, Vo, or Vehicle administration, but no consistent changes were seen at later times.

### Comparative analyses of the action of the TCF in reducing mechanical allodynia in mice and rats

We previously carried out detailed studies on the safety and pharmacokinetics of TCF in mice (*43*). To better understand the action of TCF on chronic pain, in the context of its pharmacodynamics, we investigated its effects in the mouse model of SNI and in the rat model of SNI, where we contrast TCF to various combinations of its constituents (Figure 4A shows timeline and experimental design; mice and rats were all males). To facilitate multiple comparisons, the allodynia response was followed for 10-14 days after a single injection: our data in Figs. 1 and 3 had amply established that a robust response was evident in this time period. As shown in Fig. 4B (Mice), left panel (Injured Paw), in 5-month old SNI mice (red), HPBCD+PEG showed a small reduction in allodynia, but not Vo+PEG in the injured paw by 14 days, while the TCF showed a robust reduction by day 3 that stabilized by day 10. No effect was seen with HPBCD+PEG, Vo+PEG or TCF in sham-injured mice (black). There was no effect seen with any of the three treatments in the healthy paw of SNI or Sham mice (Fig. 4B, right panel). Notably, our published pharmacokinetic studies show that in mice, within 30 minutes of injection, the TCF increased the plasma and brain exposure of Vo by 1.7 fold. TCF also slowed down removal, resulting in the elimination of Vo from the brain and blood in 4 h compared to 2 h (seen with Vo alone)(*43*). Hence, the reduction in allodynia detected on day 3 in the SNI mice, occurs well after Vo is known to be eliminated from the animal and sustained reduction of allodynia measured till day 10 is maintained in the absence of drug.

**Figure 4.**
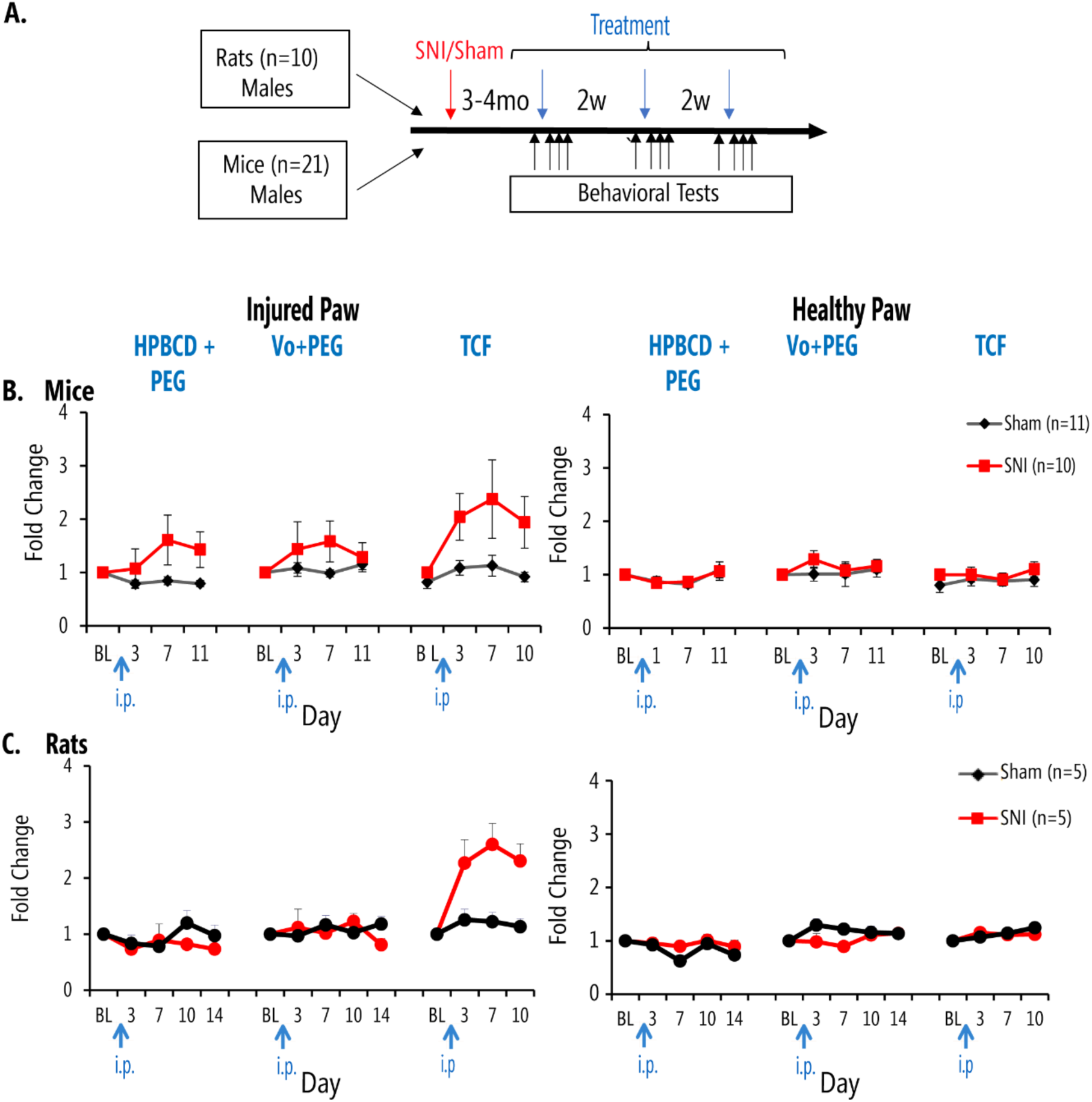
Effect of TCF in male mice and rats, with SNI and Sham injuries, relative to various combinations of constituents of TCF. **A.** Timeline and schedule of behavioral assessments (red arrow=SNI or Sham injury; blue arrows=TCF or various constituent combinations; black arrows = behavioral measures). **B**. Change in tactile allodynia in Sham (black) and SNI (red) mice, for injured (left panel) and healthy paw (right panel), for indicated TCF constituent combinations and for TCF administration. For the HPBCD+PEG drug injection, the multiple comparisons of means (Tuckey contrast) showed a significant difference between SNI and Sham mice treatment groups (p=0.0011) for the injured paw. For the TCF drug injection, there was a statistically significant difference for the injured paw (2-way, repeated-measures ANOVA 2-RM-ANOVA F_3,36_=5.208, p=0.004 for treatment*time, Supp. Table 2 Panel A, TCF). No differences for V0+PED injection or on the healthy paw **C.** Similar data as in B, for rats. Only the injured paw tactile sensitivity showed a statistically significant difference when TCF was injected (2-way, repeated-measures ANOVA 2-RM-ANOVA F_3,24_=6.649, p=0.002 for treatment*time, Supp. Table 2 Panel D, TCF).

In SNI rats (Fig. 4C, Rats, left panel), neither HPBCD+PEG nor Vo+PEG reduced allodynia in the injured paw by 14 days. However, TCF-treated animals showed measurable reduction by day 3, which improved by day 7 and was still prominent at day 10 (red). There was no effect in sham animals (black) or in the healthy paw of SNI or Sham animals (Fig. 4C; right panel). The TCF response curves in mice and rats revealed close correspondence for both the amplitude and dynamics of action of TCF in each model suggesting a shared mechanism of action between the two models.

### Effects of dose and gender on TCF action

Since the HPBCD and PEG have no effect on mechanical allodynia and the action of the TCF is to increase the available concentrations of Vo(*43*), we investigated whether changing the level of Vo in the formulation, may influence effects on chronic pain. The maximum solubility of Vo in TCF limited increasing Vo levels by twofold (100 mg/kg/day). We, therefore, compared 2xVo-TCF with 1xVo-TCF and 0.5Vo-TCF in SNI male and female rats (to also incorporate the effects of gender in these studies) (Fig. 5A experimental design and timeline). Eighteen Sprague Dawley adult rats (9 males and 9 females) were separated into two groups. In the first, six males and six females were sequentially subjected to 0.5xVo-TCF, 1xVo-TCF, and 2xVo-TCF in three treatment groups, respectively. In the second group, 3 males and 3 females were treated with vehicle as the accompanying control group.

**Figure 5.**
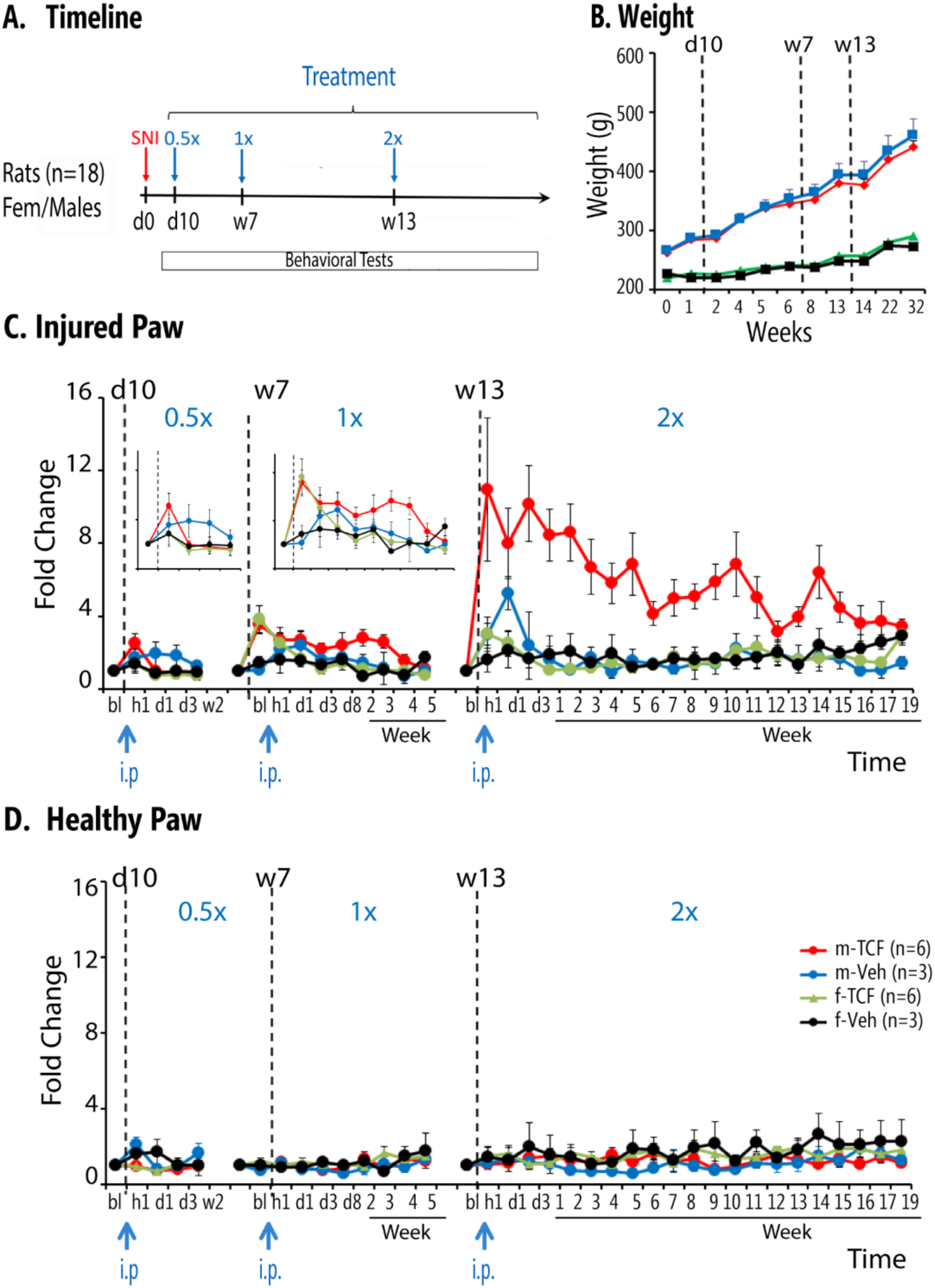
Effects of dose and gender on TCF effects in SNI rats. **A.** Timeline and schedule of treatments and behavioral assessments (red arrow=SNI injury; blue arrows=TCF or vehicle administration with increasing concentration). **B**. Body weight in male and female SNI rats exposed to TCF or vehicle throughout the duration of the study. Body weight did not differ between TCF and vehicle groups, in each sex. **C.** Tactile allodynia changes with increasing dose of TCF on the injured paw. Decreased tactile allodynia is observed mainly in male SNI. Increasing Vo concentration in TCF resulted in increased effect size and increased duration of persistence of effect (at 1.0xVo-TCF there was a drug*time effect F_9,126_ = 3.3, p < 0.002; at 2.0xVo-TCF there was a drug*sex effect F_1,14_ = 12.8, P < 0.002 and a sex*time effect F_21,294_ = 3.3, p<0.001, Supp. Table 3). **D.** Increasing Vo concentration had no effect on the healthy paw tactile sensitivity.

At day 10 after SNI, 6 adult males and 6 adult females injected with 0.5xVo and 3 males and 3 females injected with vehicle were evaluated for 2 weeks. In the 0.5Vo group, mild analgesic response was seen in males but not in females (Fig. 5C, first panel, and enlarged insert, left panel), while vehicle-treated animals showed no significant difference. After the allodynia returned to baseline, in week 7, the same 12 animals were injected with 1xVo-TCF and monitored for 5 weeks (Fig. 5C, central panel, and enlarged insert right panel) with corresponding controls injected with vehicle. A greater analgesic response to 1xVo-TCF was seen in males, which reduced to baseline by 5 weeks. A transient response was detected in females (at hour 1 that returned to baseline by day 3). After they returned to baseline, the same 12 animals were injected with 2xVo-TCF and behavior was followed for 18 weeks (Fig. 5C right panel). The amplitude of the allodynia relief response with 2xVo-TCF, increased ∼ 8 fold one hour after injection compared to 1xVo-TCF, in males only. Even after 2 months (9-10 weeks), allodynia was reduced 4-fold compared to controls. Female mice showed a mild analgesic response initially, but that fell back to baseline by days 3-4. Fig. 5D shows this treatment did not perturb the healthy paw tactile sensitivity. Raw data of this study are shown in Supp Fig. 5. Together, these data strongly support that the action of the TCF is gender specific for males, and the extent of reduction of mechanical allodynia effected is dependent on the concentration of Vo incorporated into TCF. In these animals and for 0.5xVo-TCF, 1.0xVo-TCF, and 2.0xVo-TCF, we assessed mobility and anxiety in all 4 subgroups (male/female, TCF/Veh) at baseline, hour 2 and day 11 for each treatment (Supp. Fig. 5.2). We observe small and transient perturbations (hour 2) mainly in males-TCF group and at 2.0xVo-TCF, that normalize by day 11. Cold sensitivity was also tested in these animals (acetone test) at multiple timepoints and at all 3 TCF concentrations (Supp. Fig. 5.3). We only observe transient (hour 1) decreased sensitivity only at high TCF concentrations in male-TCF group. In a previous study we have demonstrated that pain relief in SNI animals is accompanied by improved novel object recognition (*53*). Therefore, we tested novel object recognition in the last group of animals, repeatedly after 1.0xVo-TCF, and 2.0xVo-TCF (Supp. Fig. 6). We repeatedly observed object discrimination only in male SNI rats treated with TCF. The group specificity of the effect suggests that the effect is likely mediated through pain relief.

## Discussion

Our previous studies have shown that the TCF is needed to deliver a functional level of Vo into the brain, not achieved by systemic administration of Vo alone (*43*). Our present finding that the TCF is effective at reducing allodynia while Vo alone is not, also supports that TCF acts on the central nervous system. The effect on spinal cord inflammation, particularly microglia, further strengthens evidence for TCF’s action in the central nervous system. Microglial increase associated with sidedness of injury in SNI animals receiving vehicle are also consistent with prior studies showing that they are associated with spinal cord inflammation (*52*)). TCF reduced microglial levels to baseline. It also decreased levels of astrocytes (detected by GFAP), suggesting one mechanism of action of the TCF may be through the reduction of glia. However other responses of the central nervous and peripheral nervous system may also play a role and need to be further investigated. The outcomes of a large set of phenotypic assays suggest that TCF may have a long-lasting analgesic effect on mechanical allodynia mediated by large fibers with less involvement on smaller fibers and no side effects on tactile sensitivity for uninjured paw, weight, anxiety, and mobility.

When a single dose of TCF is systemically administered in mice, Vo levels rise in plasma and brain to peak at 30 min and then rapidly fall, and then rapidly eliminated from the brain and body in 4h (*43*). However, the reduction in allodynia is sustained for 3-4 weeks, suggesting it is sustained by mechanisms of action for 25-30 days, well after elimination of Vo. This is consistent Vo/HDACi-mediated long-term epigenetic processes playing an important role in reducing allodynia. Comparison of dynamics of reduction in allodynia in rat and mouse models suggests a similar mechanism of action in both species. Doubling the amount of vorinostat in TCF greatly boosted the amplitude and duration of reduced allodynia, suggesting initial levels of target modification may impact duration of efficacy. With the 2x dose, after four months, mechanical allodynia is eventually re-established. In future studies, it may be interesting to investigate dose-dependent epigenetic modification (e.g. histone acetylation) and whether their long-term waning is commensurate with the increase in allodynia. Since Vo is known to act broadly across HDAC1 and HDAC2, indirect effects (beyond histone acetylation) may also play a role.

In the present studies, the observed effects of TCF were mainly for tactile sensitivity with small effects on cold. It is possible that higher concentrations of TCF may affect cold more robustly, as suggested by the 2x dose effects. We did not test heat as this modality is minimally disturbed in SNI. Other neuropathic models, like chronic constriction injury (CCI) to the sciatic nerve, have a larger heat hyperalgesia that could be tested for TCF effects. Testing TCF effects on other rodent models of pain would also be informative, such as acute or long-term peripheral inflammatory models, and cancer chemotherapy pain models (*54*). We did not test the extent to which the affective component of SNI neuropathic pain was being modulated by TCF. Such experiments become complicated due mainly to the long-lasting effects of TCF. Our evidence that social recognition improved with TCF only in male SNI animals, itself suggests that TCF is also reducing affective burden of SNI. It is possible that TCF could ameliorate acute pain conditions too. But this is not a domain where the mechanism would have real utility. In contrast, chronic pain conditions like back pain, osteoarthritis and fibromyalgia are all domains where available treatments, especially pharmacotherapy, remains inadequate commonly leading to unacceptable side effects.

In SNI rats, a single dose of TCF shows comparable efficacy to weekly administration, consistent with the idea that epigenetic effects from a single dose last ∼3-4 weeks. However, doubling Vo levels in TCF increases both amplitude and duration of efficacy to ∼ 3 months. This is a unique property of TCF, no current treatments of chronic pain show such effects. In sum, through pharmacological modulation of Vo, the TCF enables single-dose effectiveness with extended action and reduces long-term HDACi dosage. A treatment with long-duration efficacy decreases exposing the patient to iatrogenic effects. Thus, TCF has the potential to revolutionize management of chronic pain.

## Acknowledgements.

The study was in part supported by NIH NIDA P50 DA044121 Director A.V.A. and in part by the University of Notre Dame through the Parsons-Quinn Endowment, (M.S.A) and sabbatical leave to K.H. Author contributions: M.V.C. maintained all rat and mouse colonies, helped conceive and design the studies, performed all the injections as well as live animal assessments in the rat and mouse models model as well as organ harvest, analyzed the data in Figures 1,3-5 and Supplementary figures and wrote the paper. M.S.A. helped conceive and design the study, prepared TCF and vehicle formulations, undertook RT-PCR experiments and quantitative analyses of immunolocalization studies, assembly and analyses of data in Fig. 2, and edited the paper. K.H. and A.V.A. conceived and designed the study and its experiments, analyzed the data and wrote the paper.

## Supplementary Material

**Supplementary Figure 1.**
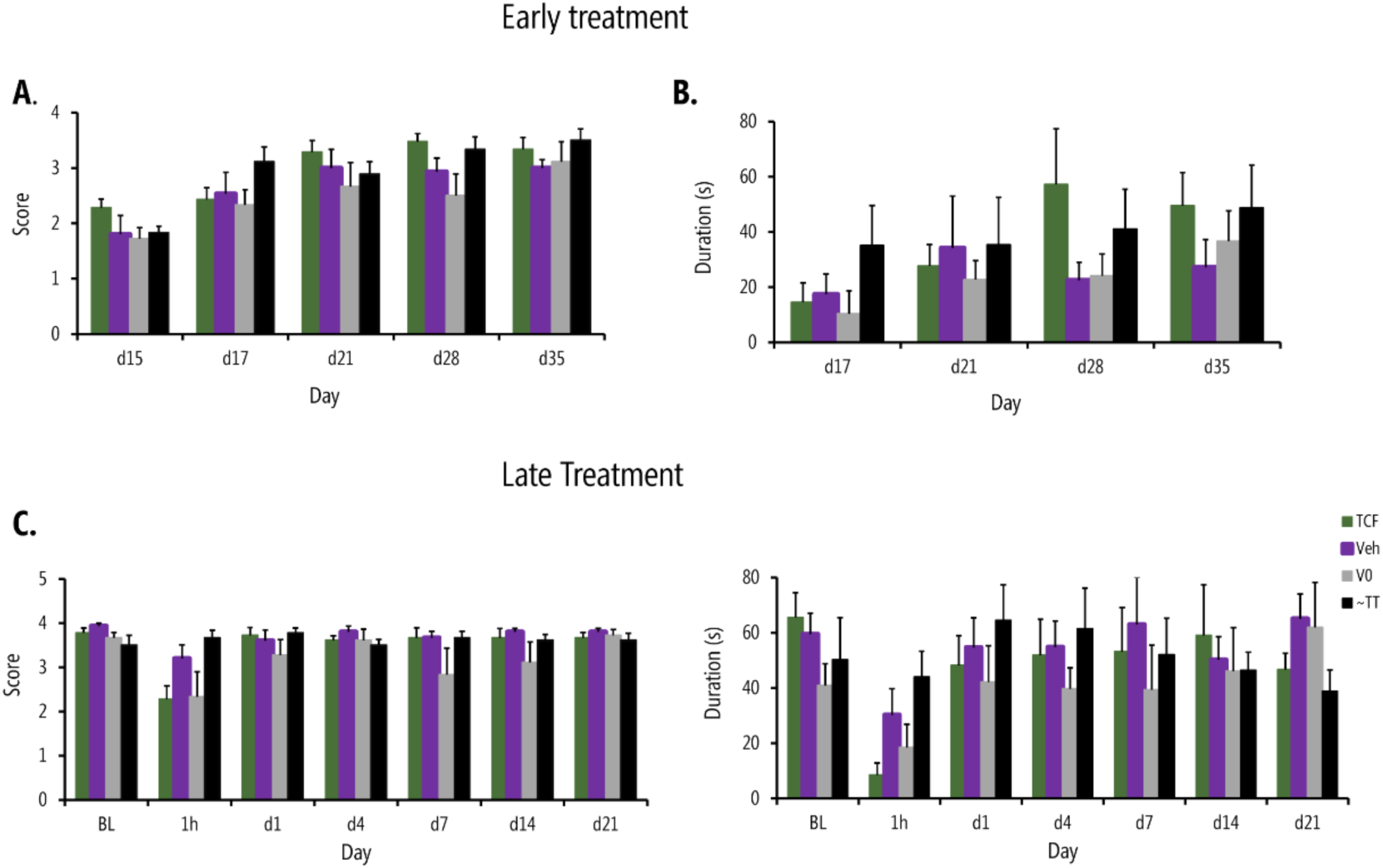
Cold sensitivity tested using acetone test. Rats in the group described in Figure 3 (main manuscript) were repeatedly tested for withdrawal signs to acetone applied to SNI injured paw. 24 male SNI rats: 7 treated with TCF, 6 vehicle, 5 Vo, and 6 no treatment. Average withdrawal score (**A, C**) and average duration of paw withdrawal (**B, D**) are shown, for early (**A, B**) and late (**C, D**) treatments. Overall, we observe minimal changes in cold sensitivity, mainly within the first hour after either TCF or Vo, which transiently seem to diminish cold sensitivity.

**Supplementary Figure 2.**
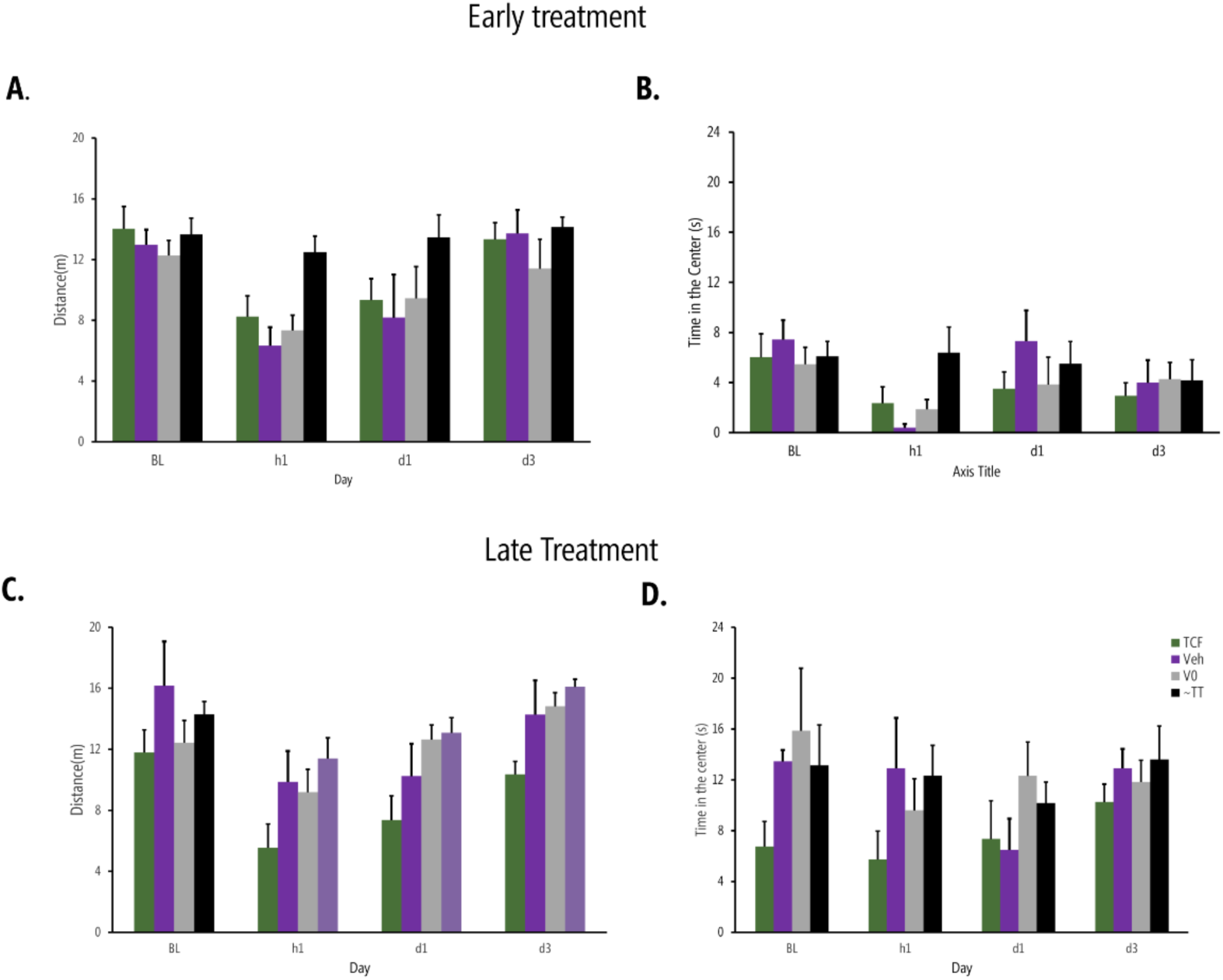
Mobility and anxiety were assessed in open field task. Rats in the group described in Figure 3 (main manuscript) were repeatedly tested. 24 male SNI rats: 7 treated with TCF, 6 vehicle, 5 Vo, and 6 no treatment. Distance traveled in 5 minutes (mobility) (**A, C**) and time in the center (anxiety assay) (**B, D**) are shown, for early (**A, B**) and late (**C, D**) treatments. Overall, we observe minimal changes in these outcomes. The data shows large variability and possible differences between the groups all dissipating by days 1-3 after treatments.

**Supplementary Figure 3.**
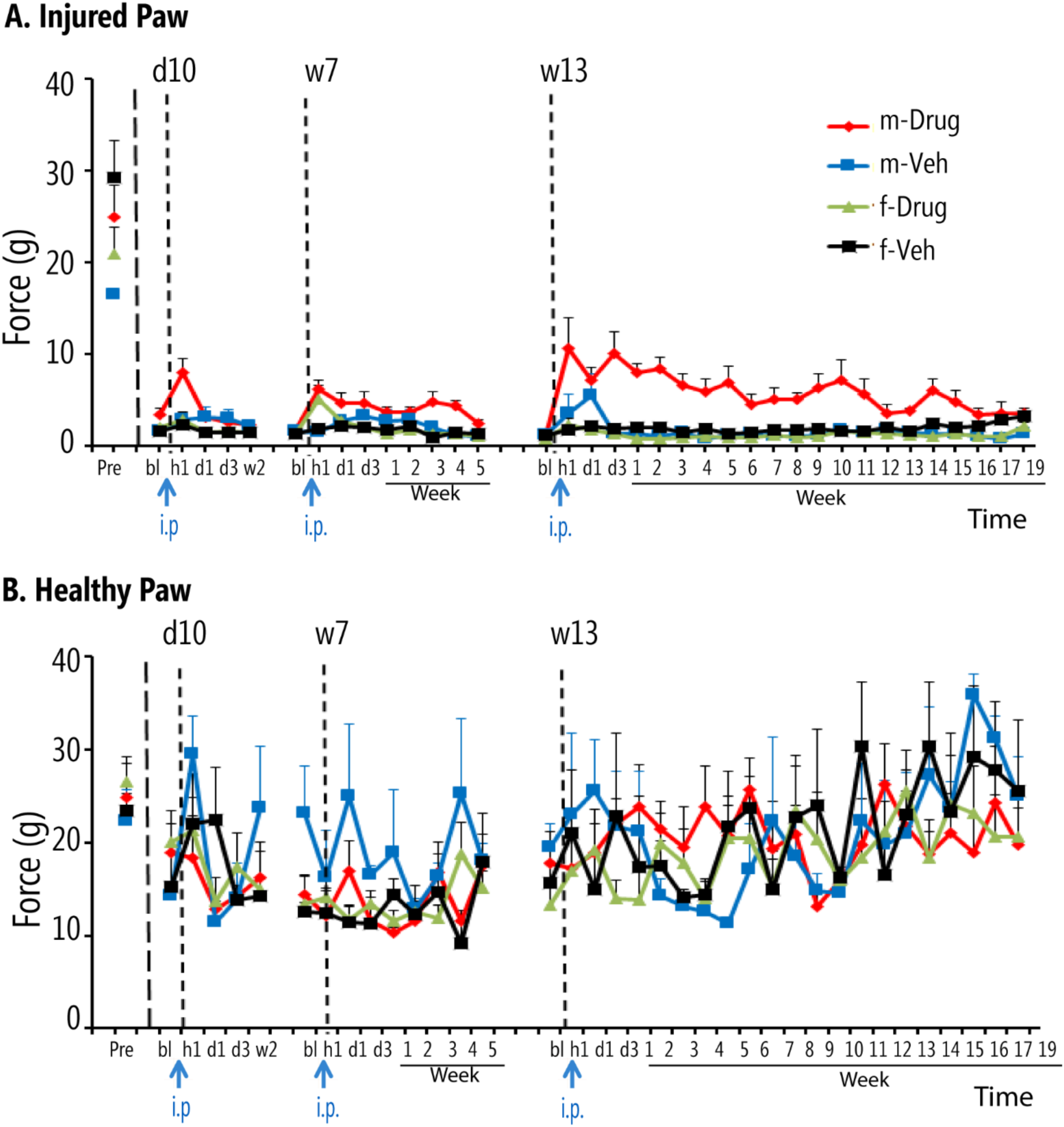
Same data as in the main manuscript figure 5, shown in absolute scale.

**Supplementary Figure 4.**
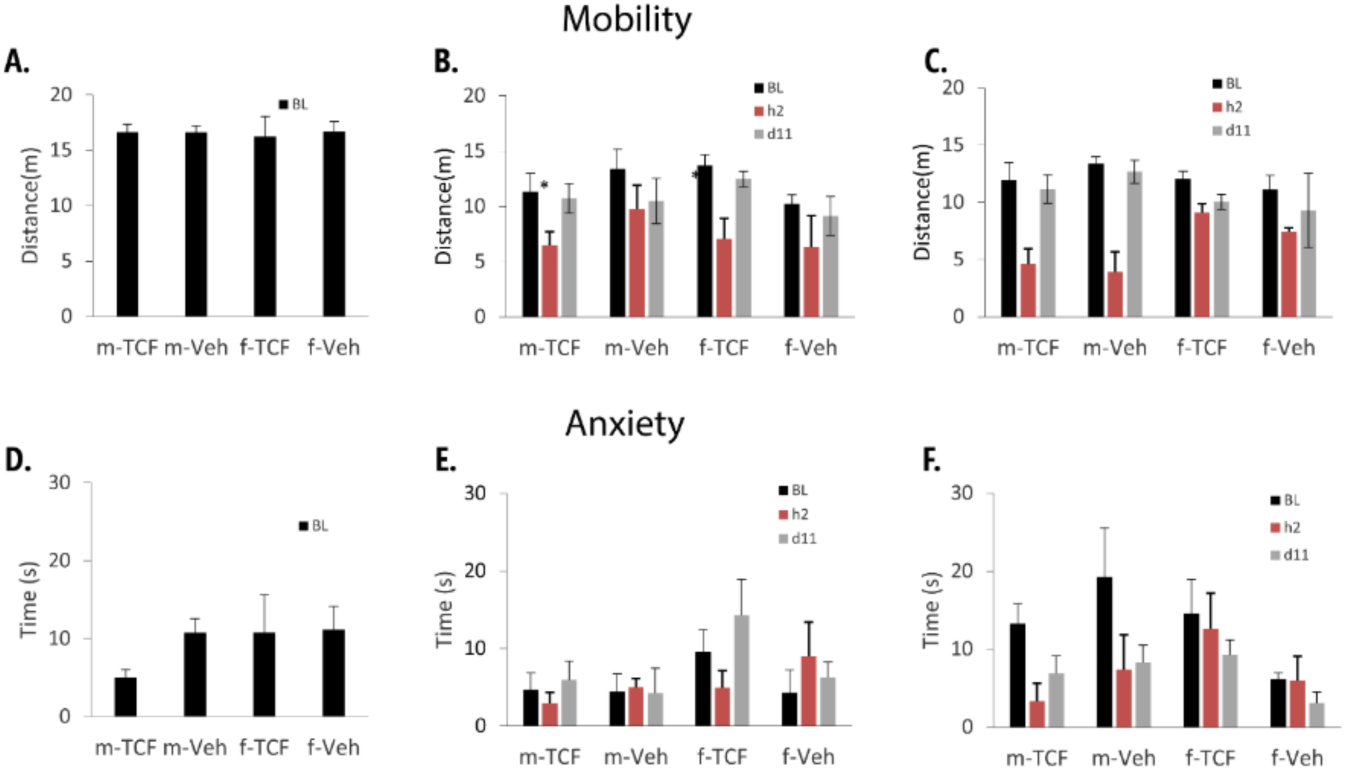
Mobility and anxiety were assessed prior to treatment (BL), 2 hours after treatment (h2), and 11 days after the start of treatment (d11) for all four animal groups described in the main manuscript figure 5 and supplementary figure 3. **A-C** are distance traveled in 5 minutes open field, **D-F** are time in center. Data are shown for 1xVo-TCF and 2xVo-TCF. We observe no long-term mobility changes, but perhaps some increased anxiety (less time spent in center) for 2xVo-TCF in males treated with TCF and vehicle.

**Supplementary figure 5:**
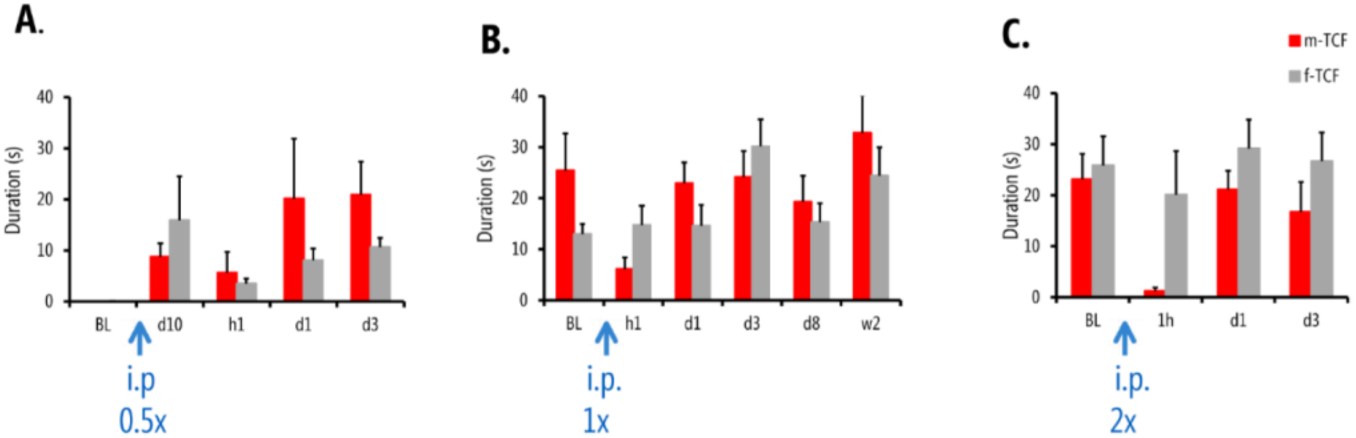
Cold sensitivity (acetone test) tested in male and female TCF treated SNI rats, for increasing doses of Vo in TCF. We observe decreased cold sensitivity at hour 1 in males treated with TCF (m-TCF) for 1xVo-TCF, and more sustained small decrease in cold sensitivity (day 3) in m-TCF for 2xVo-TCF.

**Supplementary Figure 6:**
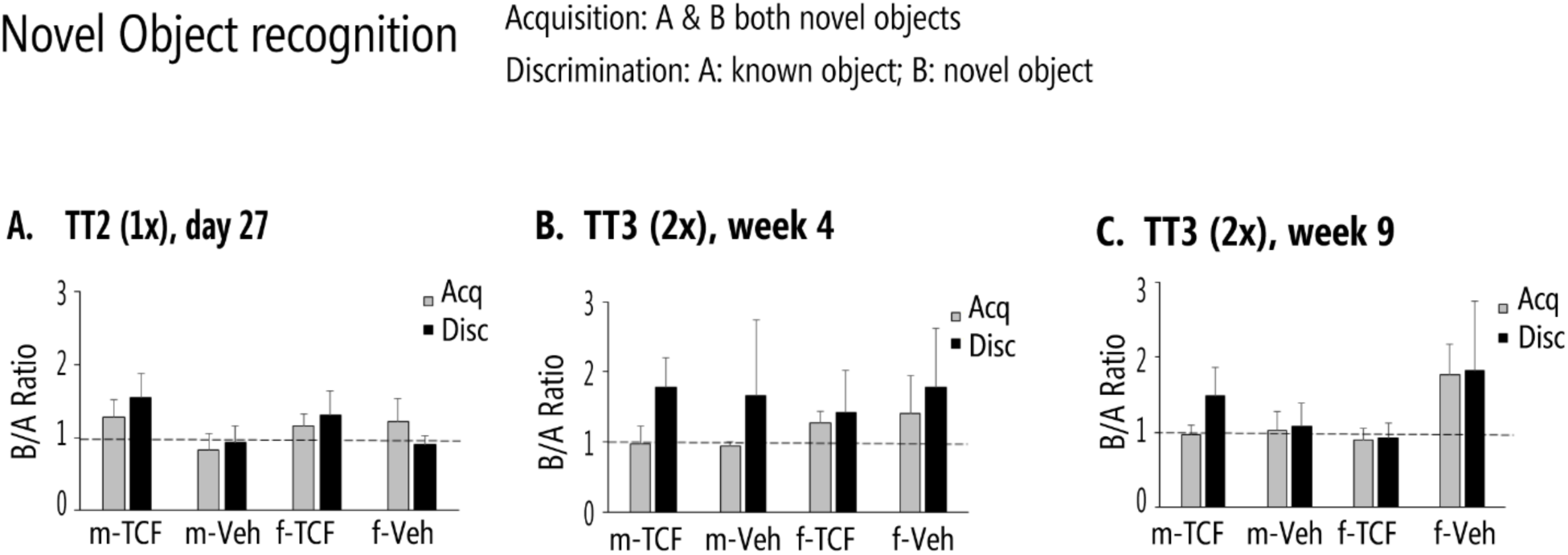
Novel object recognition improves only in male TCF, especially at 2.0xVo-TCF doses, both at week 4 and week 9.

**Supplementary Table 1.**
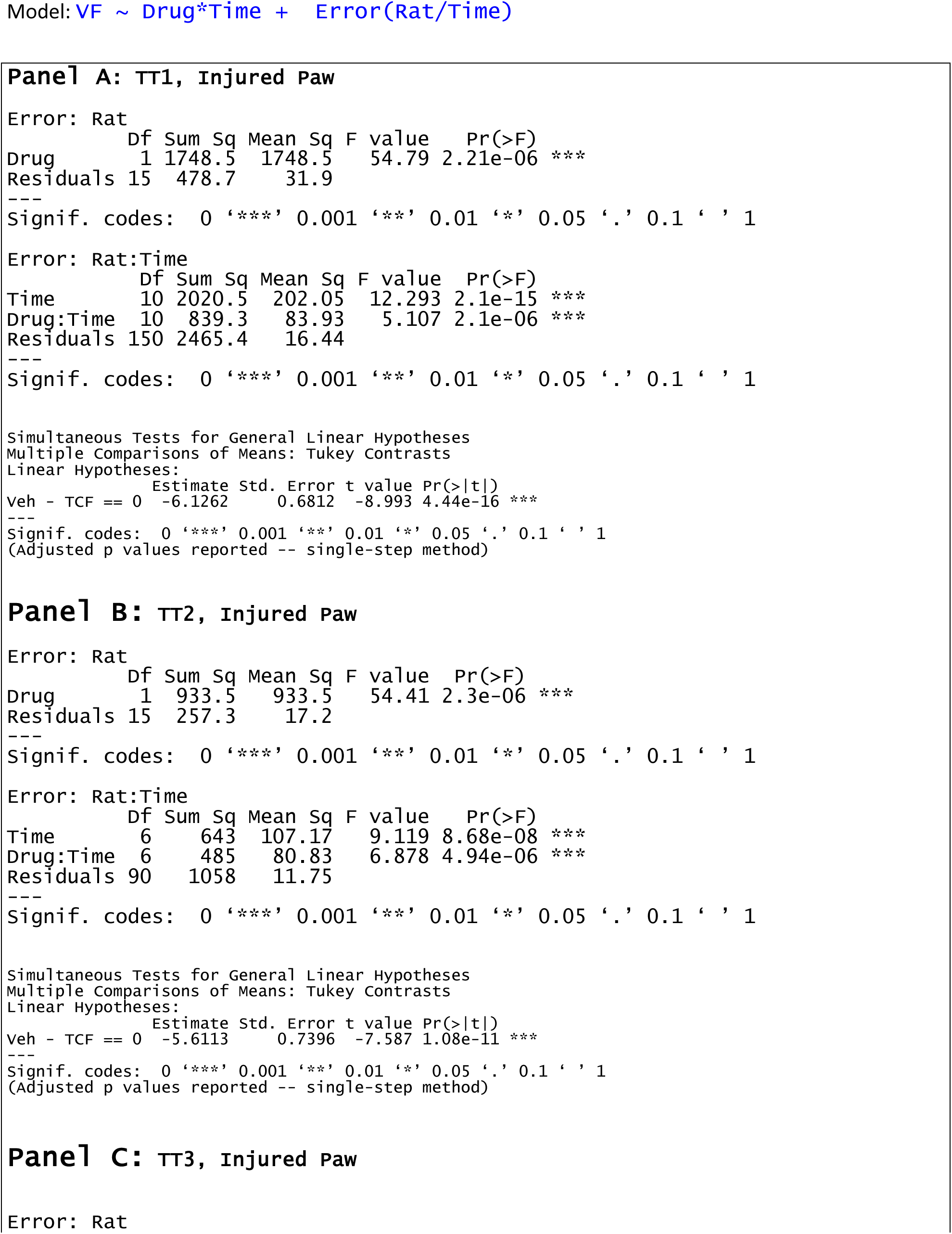
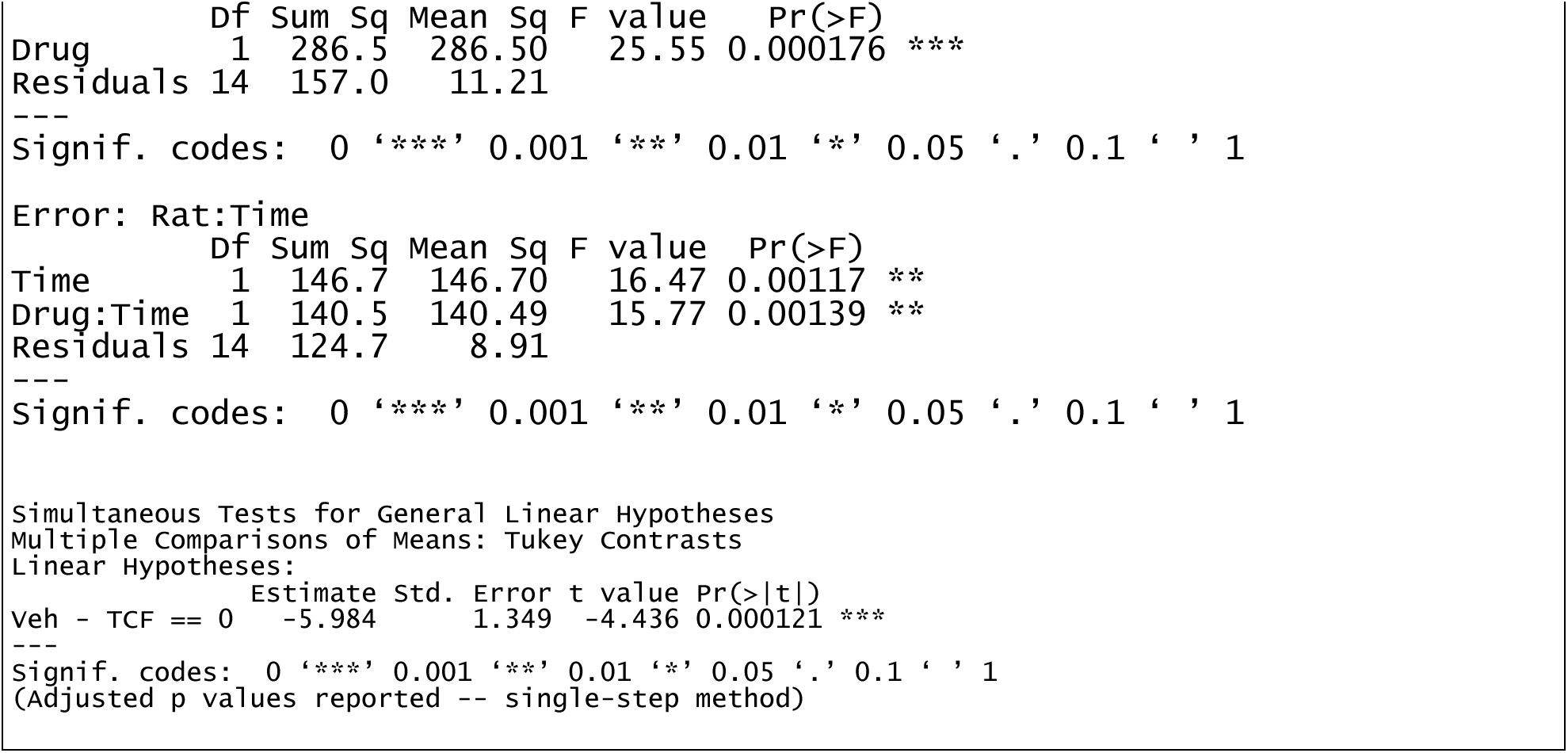
Statistical results for main Figure 1 data.

|  | Df | Sum Sq | Mean Sq | F value | Pr(>F) |
| --- | --- | --- | --- | --- | --- |
| Drug | 1 | 1748.5 | 1748.5 | 54.79 | 2.21e-06 *** |
| Residuals | 15 | 478.7 | 31.9 |  |  |

|  | Df | Sum Sq | Mean Sq | F value | Pr(>F) |
| --- | --- | --- | --- | --- | --- |
| Time | 10 | 2020.5 | 202.05 | 12.293 | 2.1e-15 *** |
| Drug:Time | 10 | 839.3 | 83.93 | 5.107 | 2.1e-06 *** |
| Residuals | 150 | 2465.4 | 16.44 |  |  |

|  | Estimate | Std. Error | t value | Pr(>\|t\|) |
| --- | --- | --- | --- | --- |
| Veh - TCF == 0 | -6.1262 | 0.6812 | -8.993 | 4.44e-16 *** |

|  | Df | Sum Sq | Mean Sq | F value | Pr(>F) |
| --- | --- | --- | --- | --- | --- |
| Drug | 1 | 933.5 | 933.5 | 54.41 | 2.3e-06 *** |
| Residuals | 15 | 257.3 | 17.2 |  |  |

|  | Df | Sum Sq | Mean Sq | F value | Pr(>F) |
| --- | --- | --- | --- | --- | --- |
| Time | 6 | 643 | 107.17 | 9.119 | 8.68e-08 *** |
| Drug:Time | 6 | 485 | 80.83 | 6.878 | 4.94e-06 *** |
| Residuals | 90 | 1058 | 11.75 |  |  |

|  | Estimate | Std. Error | t value | Pr(>\|t\|) |
| --- | --- | --- | --- | --- |
| Veh - TCF == 0 | -5.6113 | 0.7396 | -7.587 | 1.08e-11 *** |
| Drug | 1 | 286.5 | 286.50 | 25.55 | 0.000176 *** |
| Residuals | 14 | 157.0 | 11.21 |  |  |

|  | Df | Sum Sq | Mean Sq | F value | Pr(>F) |
| --- | --- | --- | --- | --- | --- |
| Time | 1 | 146.7 | 146.70 | 16.47 | 0.00117 ** |
| Drug:Time | 1 | 140.5 | 140.49 | 15.77 | 0.00139 ** |
| Residuals | 14 | 124.7 | 8.91 |  |  |

|  | Estimate | Std. Error | t value | Pr(>\|t\|) |
| --- | --- | --- | --- | --- |
| veh - TCF == 0 | -5.984 | 1.349 | -4.436 | 0.000121 *** |

--- Signif. codes: 0 '\*\*\*' 0.001 '\*\*' 0.01 '\*' 0.05 '.' 0.1 ' ' 1 (Adjusted p values reported -- single-step method)

**Supplementary Table 2.**
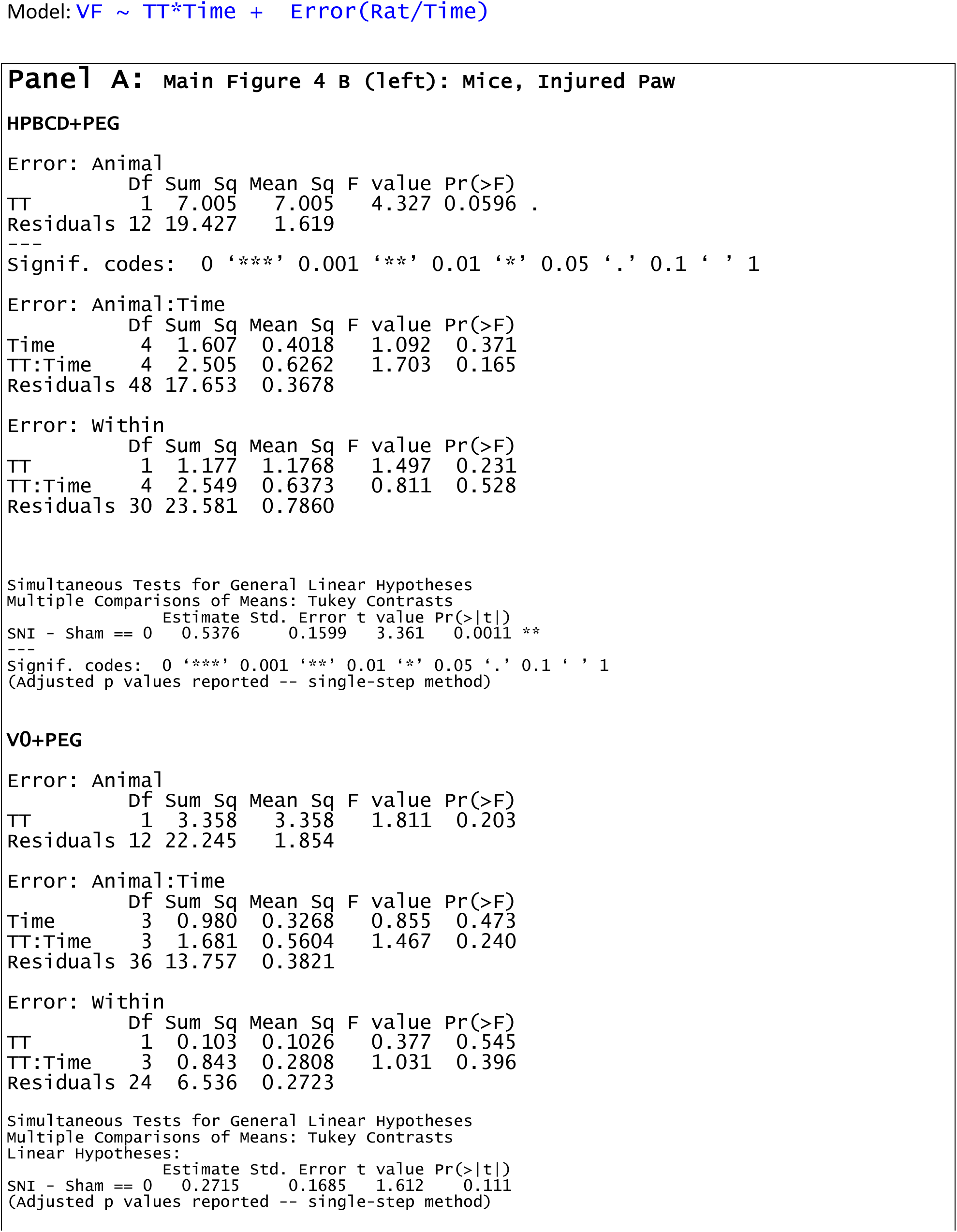
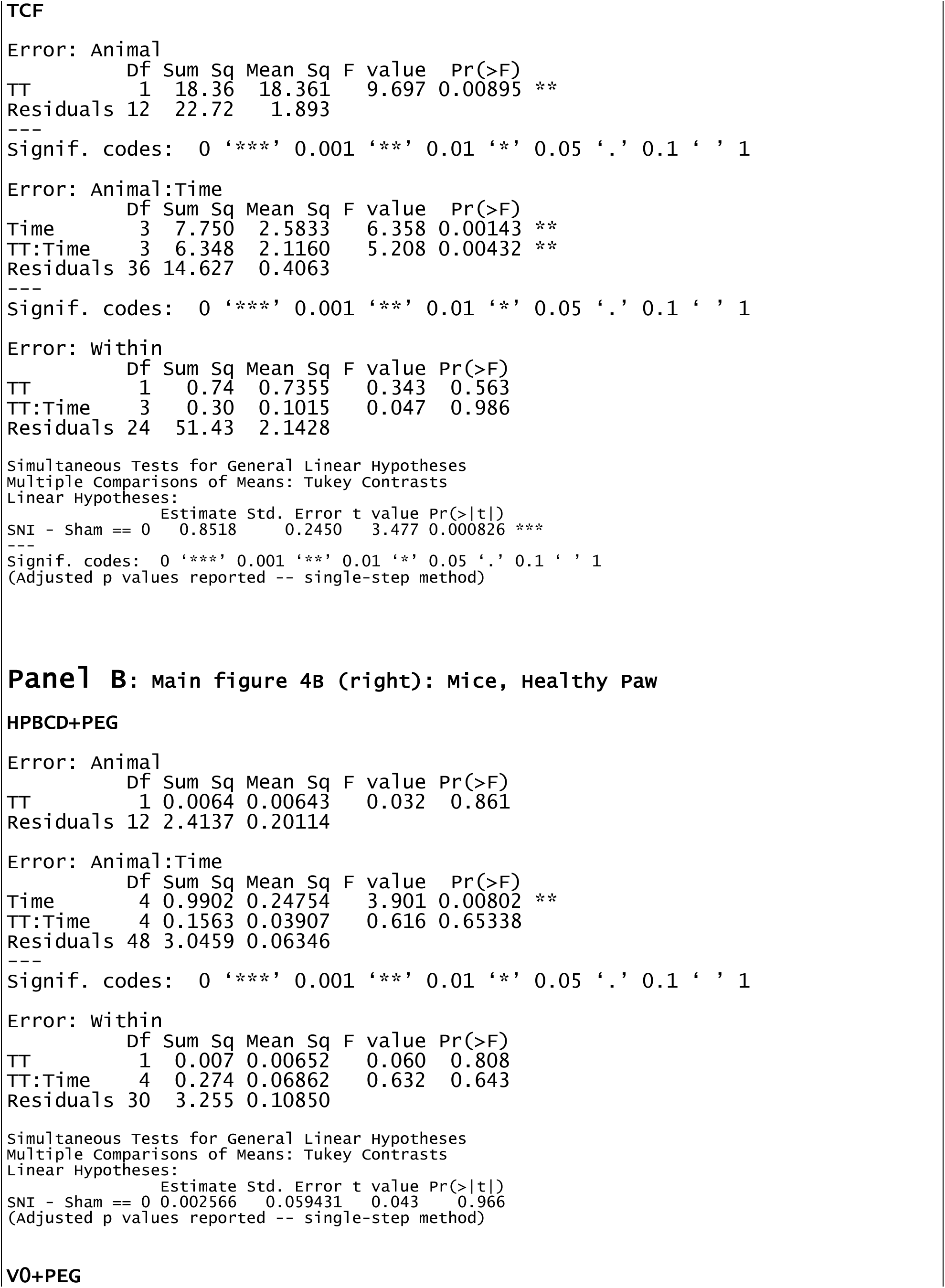
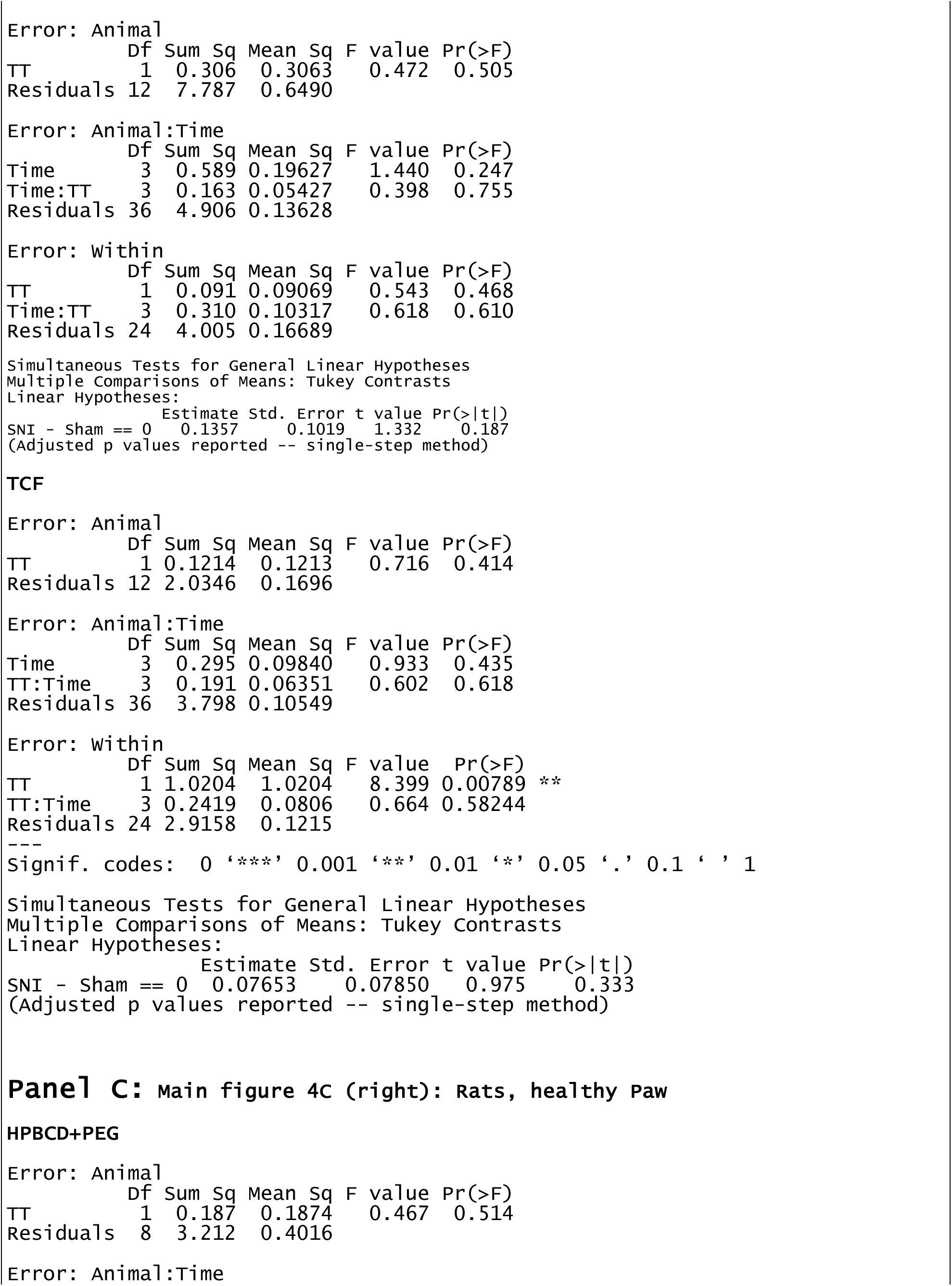
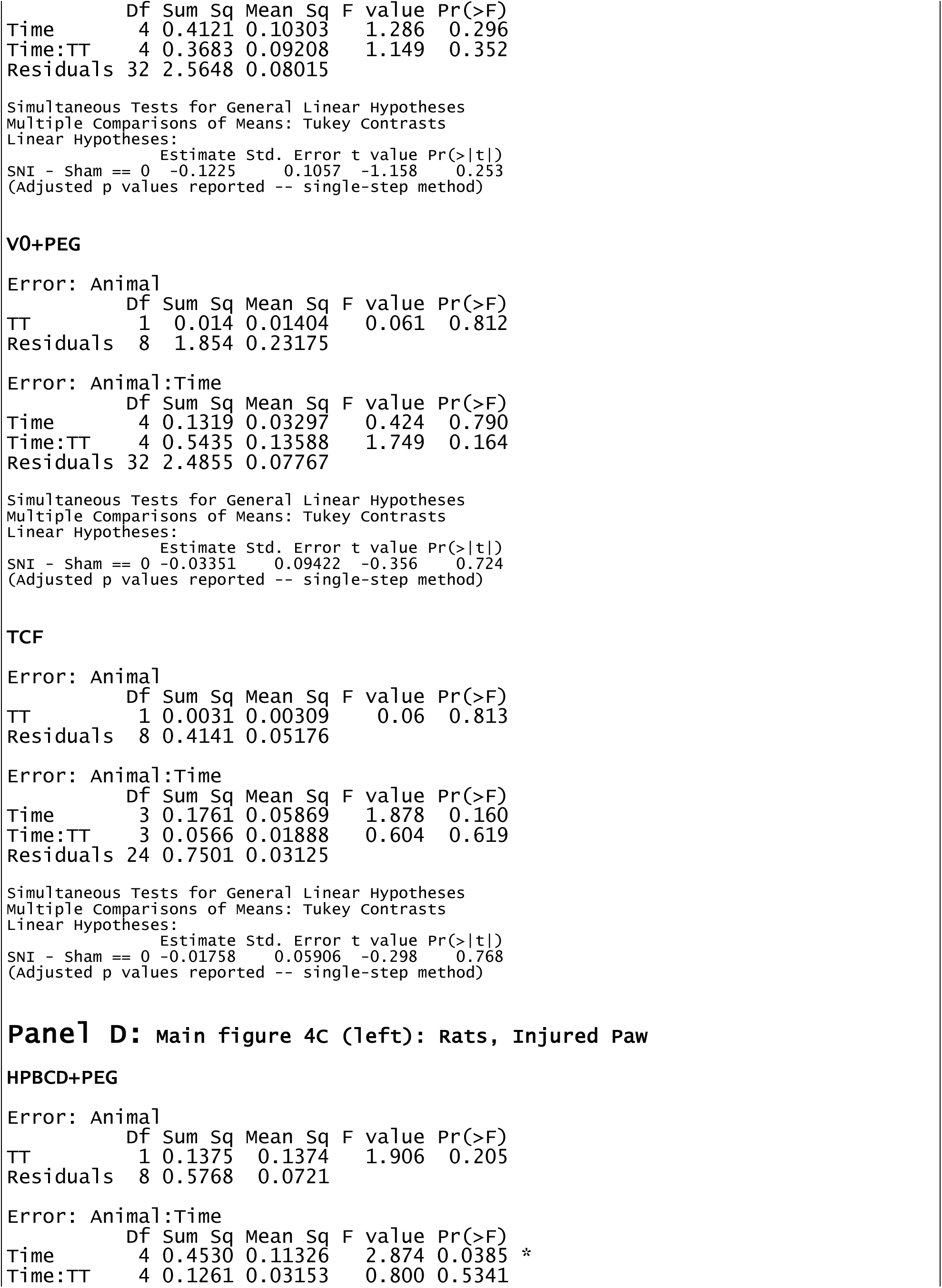
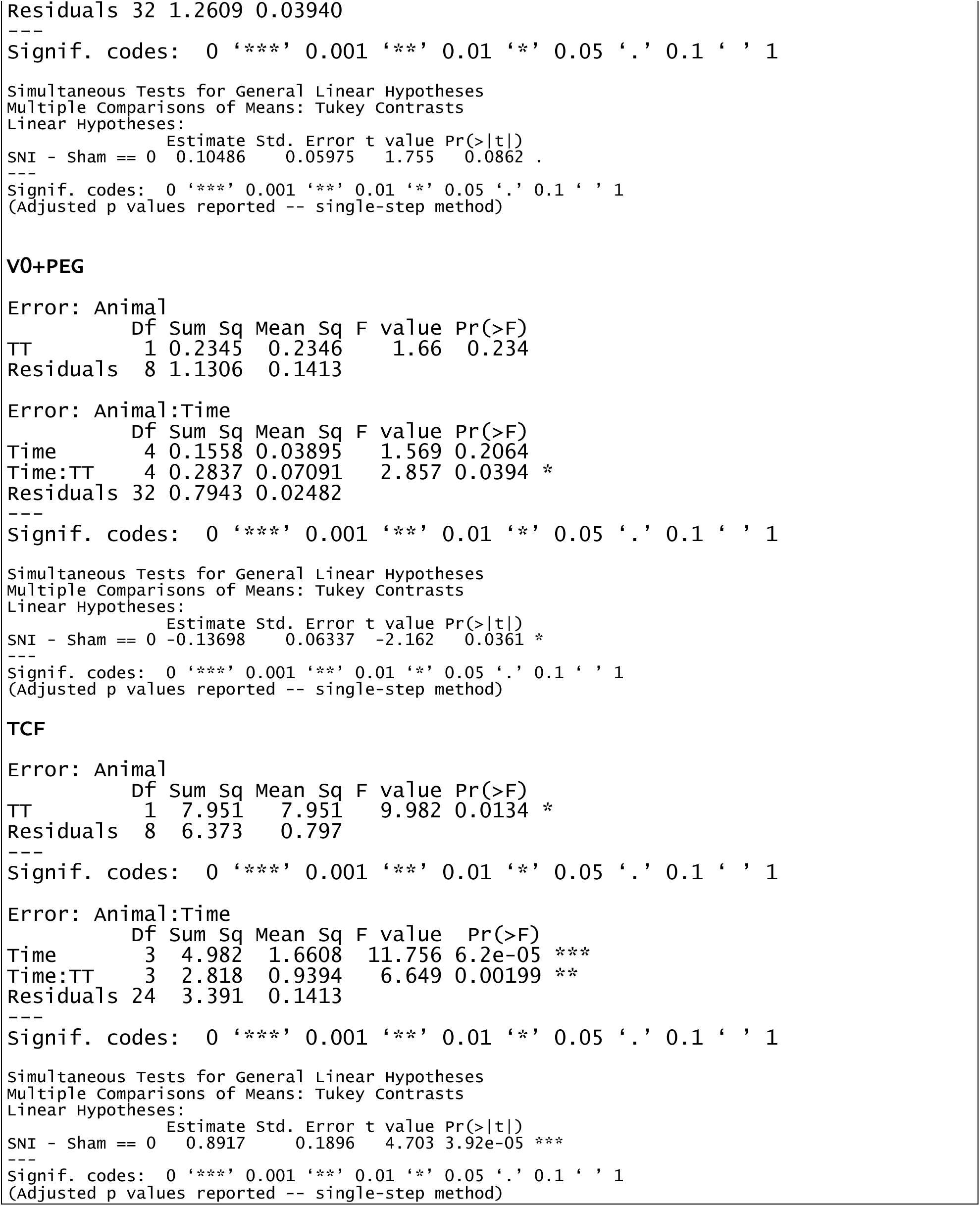
Statistical results for main Figure 4 data.

**Supplementary Table 3.** Statistical results for main Figure 5 data.

|  | Df | Sum Sq | Mean Sq | F value | Pr(>F) |
| --- | --- | --- | --- | --- | --- |
| Drug | 1 | 0.915 | 0.9146 | 0.834 | 0.377 |
| Sex | 1 | 2.983 | 2.9825 | 2.720 | 0.121 |
| Drug:Sex | 1 | 0.257 | 0.2572 | 0.235 | 0.636 |
| Residuals | 14 | 15.350 | 1.0964 |  |  |

Error: Rat:Time
|  | Df | Sum Sq | Mean Sq | F value | Pr(>F) |
| --- | --- | --- | --- | --- | --- |
| Time | 4 | 10.904 | 2.7259 | 10.633 | 1.76e-06 *** |
| Drug:Time | 4 | 2.644 | 0.6611 | 2.579 | 0.0471 * |
| Sex:Time | 4 | 2.037 | 0.5092 | 1.986 | 0.1091 |
| Drug:Sex:Time | 4 | 1.746 | 0.4364 | 1.702 | 0.1623 |
| Residuals | 56 | 14.356 | 0.2564 |  |  |

|  | Estimate | Std. Error | t value | Pr(>\|t\|) |
| --- | --- | --- | --- | --- |
| Veh - TCF == 0 | 0.2138 | 0.1481 | 1.444 | 0.152 |

|  | Df | Sum Sq | Mean Sq | F value | Pr(>F) |
| --- | --- | --- | --- | --- | --- |
| Drug | 1 | 11.78 | 11.780 | 3.857 | 0.0697 . |
| Sex | 1 | 11.90 | 11.905 | 3.898 | 0.0684 . |
| Drug:Sex | 1 | 3.42 | 3.421 | 1.120 | 0.3078 |
| Residuals | 14 | 42.76 | 3.054 |  |  |

|  | Df | Sum Sq | Mean Sq | F value | Pr(>F) |
| --- | --- | --- | --- | --- | --- |
| Time | 9 | 59.27 | 6.585 | 8.883 | 2.77e-10 *** |
| Drug:Time | 9 | 21.72 | 2.413 | 3.255 | 0.00137 ** |
| Sex:Time | 9 | 12.07 | 1.341 | 1.809 | 0.07266 . |
| Drug:Sex:Time | 9 | 2.92 | 0.325 | 0.438 | 0.91225 |
| Residuals | 126 | 93.41 | 0.741 |  |  |

|  | Estimate | Std. Error | t value | Pr(>\|t\|) |
| --- | --- | --- | --- | --- |
| Veh - TCF == 0 | -0.5427 | 0.1620 | -3.35 | 0.000997 *** |

|  | Df | Sum Sq | Mean Sq | F value | Pr(>F) |
| --- | --- | --- | --- | --- | --- |
| Drug | 1 | 339.0 | 339.0 | 11.40 | 0.004516 ** |
| Sex | 1 | 718.1 | 718.1 | 24.15 | 0.000228 *** |

|  | Df | Sum Sq | Mean Sq | F value | Pr(>F) |  |
| --- | --- | --- | --- | --- | --- | --- |
| Time | 21 | 355.6 | 16.932 | 4.083 | 2.02e-08 | *** |
| Drug:Time | 21 | 102.6 | 4.886 | 1.178 | 0.269 |  |
| Sex:Time | 21 | 289.7 | 13.797 | 3.327 | 2.46e-06 | *** |
| Drug:Sex:Time | 21 | 100.0 | 4.761 | 1.148 | 0.297 |  |
| Residuals | 294 | 1219.2 | 4.147 |  |  |  |

|  | Estimate | Std. Error | t value | Pr(>\|t\|) |  |
| --- | --- | --- | --- | --- | --- |
| Veh - TCF == 0 | -1.9628 | 0.2767 | -7.093 | 6.68e-12 | *** |

``` Signif. codes: 0 '***' 0.001 '**' 0.01 '*' 0.05 '.' 0.1 ' ' 1 (Adjusted p values reported -- single-step method) ```

